# Cell-intrinsic platinum response and associated genetic and gene expression signatures in ovarian cancer cell lines and isogenic models

**DOI:** 10.1101/2024.07.26.605381

**Authors:** Kristin M. Adams, Jae-Rim Wendt, Josie Wood, Sydney Olson, Ryan Moreno, Zhongmou Jin, Srihari Gopalan, Jessica D. Lang

## Abstract

Ovarian cancers are still largely treated with platinum-based chemotherapy as the standard of care, yet few biomarkers of clinical response have had an impact on clinical decision making as of yet. Two particular challenges faced in mechanistically deciphering platinum responsiveness in ovarian cancer have been the suitability of cell line models for ovarian cancer subtypes and the availability of information on comparatively how sensitive ovarian cancer cell lines are to platinum. We performed one of the most comprehensive profiles to date on 36 ovarian cancer cell lines across over seven subtypes and integrated drug response and multiomic data to improve on our understanding of the best cell line models for platinum responsiveness in ovarian cancer. RNA-seq analysis of the 36 cell lines in a single batch experiment largely conforms with the currently accepted subtyping of ovarian cancers, further supporting other studies that have reclassified cell lines and demonstrate that commonly used cell lines are poor models of high-grade serous ovarian carcinoma. We performed drug dose response assays in the 32 of these cell lines for cisplatin and carboplatin, providing a quantitative database of IC_50_s for these drugs. Our results demonstrate that cell lines largely fall either well above or below the equivalent dose of the clinical maximally achievable dose (C_max_) of each compound, allowing designation of cell lines as sensitive or resistant. We performed differential expression analysis for high-grade serous ovarian carcinoma cell lines to identify gene expression correlating with platinum-response. Further, we generated two platinum-resistant derivatives each for OVCAR3 and OVCAR4, as well as leveraged clinically-resistant PEO1/PEO4/PEO6 and PEA1/PEA2 isogenic models to perform differential expression analysis for seven total isogenic pairs of platinum resistant cell lines. While gene expression changes overall were heterogeneous and vast, common themes were innate immunity/STAT activation, epithelial to mesenchymal transition and stemness, and platinum influx/efflux regulators. In addition to gene expression analyses, we performed copy number signature analysis and orthogonal measures of homologous recombination deficiency (HRD) scar scores and copy number burden, which is the first report to our knowledge applying field-standard copy number signatures to ovarian cancer cell lines. We also examined markers and functional readouts of stemness that revealed that cell lines are poor models for examination of stemness contributions to platinum resistance, likely pointing to the fact that this is a transient state. Overall this study serves as a resource to determine the best cell lines to utilize for ovarian cancer research on certain subtypes and platinum response studies, as well as sparks new hypotheses for future study in ovarian cancer.

## Introduction

Ovarian cancers, of which high-grade serous ovarian carcinoma (HGSOC) is the most common subtype, account for 5% of all cancer deaths in females in the United States^1^. Current standard of care for HGSOC is platinum-based chemotherapy (carboplatin, cisplatin + taxane)^2^, but over half of patients are diagnosed with Stage IV HGSOC and face a dismal 31.5% five-year survival rate^3^. Overall, the progression free interval for first-line OvCa is ∼18 months, and around 80% of patients are considered sensitive to platinum-based chemotherapy^4^. However, patients initially sensitive to platinum-based chemotherapy display decreased progression-free intervals and response rates on platinum with each subsequent recurrence^5^. OvCa patients often undergo prolonged periods of remission following initial response to surgery and platinum-based chemotherapy before ultimately recurring, frequently with chemoresistance^6^. Currently, the only way to determine if a patient is responsive to platinum-based chemotherapy is to treat them and wait up to six months to determine response.

Platinum-based chemotherapies work by binding to DNA to form inter- and intrastrand cisplatin-DNA adducts that block DNA synthesis, leading to cell cycle arrest and induction of apoptosis^7^. These DNA adducts are repaired by a variety of DNA repair mechanisms. These adducts are predominantly cleared by nucleotide excision repair (NER), which involves the nuclease heterodimer ERCC1-XPF^8^. Increased expression of ERCC1 has been identified as a predictor of cisplatin resistance^9^. Other DNA repair mechanisms involved in detecting and repairing cisplatin-DNA adducts include mismatch repair (MMR), non-homologous end joining (NHEJ), and homologous recombination repair (HRR)^10^. Deficiencies in these DNA repair pathways have also been implicated in predicting response to platinum therapies^11^. The proteins BRCA1 and BRCA2 are major proteins involved in maintaining the integrity of HRR processes^12^, and epithelial ovarian cancers with deficiencies in BRCA1/2 show increased sensitivity to platinum therapy^13^. Cisplatin is known to cause an accumulation of reactive oxygen species (ROS), causing oxidative stress which, in addition to genotoxic stress, can trigger induction of apoptosis^14^. Another predominant mechanism of multidrug resistance, which is applicable to platinum resistance as well, is the upregulation of drug efflux or inhibition of drug influx, effectively decreasing the intracellular concentrations of the active molecule. In addition, a host of other oncogenic signaling pathways have been implicated in resistance to platinum, which are reviewed elsewhere^15^.

It is important that cell line models are accurately classified and reflect the molecular characteristics of their assigned subtypes. The cell lines SKOV3 and A2780 are the two most cited cell lines in OvCa studies, and while much of OvCa research is focused on HGSOC, these two subtypes have been determined to be poor models of HGSOC and are likely ovarian clear cell carcinoma (OCCC) and endometrioid ovarian carcinoma (EOC), respectively^16,17^. As classifications of ovarian cancer have evolved over the last two decades, multiple recent studies have sought to accurately classify commercially available OvCa cell lines based on their molecular characteristics^16–19^. However, there remains an unmet need in the field for a robust review of OvCa cell lines and their sensitivities to platinum therapies and robust model systems in which to examine naturally developed mechanisms of platinum resistance. Our study sought to define the platinum sensitivity of 36 OvCa cell lines and their corresponding genetic and gene expression signatures. Further, we leveraged seven isogenic platinum-resistant cell line pairs derived from four initially platinum sensitive counterparts to more robustly determine mechanisms. Our results indicate that common themes of platinum resistance are predominated by innate immunity/STAT activation, epithelial to mesenchymal transition and stemness, and platinum influx/efflux regulators. This work serves as a resource to the ovarian cancer community for more appropriate and well-defined platinum-responsiveness across multiple cell types in the aim that the field of platinum-responsiveness in ovarian cancer can be accelerated by using more appropriate model systems.

## Methods

### Cell culture

Cell lines used in this study and the media preparations used for culture are described in **Supplemental Table 1**. All cell lines were cultured in a humidified 37°C incubator with 5% CO_2_. All culture medium was supplemented with 1% penicillin/streptomycin (Gibco). Cell line authentication was performed through the University of Wisconsin-Madison’s Translational Research Initiatives in Pathology (TRIP) Lab. Mycoplasma testing was performed using the MycoAlert^TM^ Mycoplasma Detection Kit (Lonza).

### Drug dose response assay

Cells were plated in 50µL of the appropriate culture medium in a white opaque-walled 96-well tissue culture plate (Corning), and the number of cells plated per well for each cell line is detailed in **Supplemental Table 1**. The cells were allowed to attach for 24 hours before cisplatin (Cayman Chemical Company, cat. 13119) treatment, when the final volume in each well was brought to 100µL. Each cell line was tested at 10-doses that ranged between 400uM to 60nM depending on the sensitivity of the cell line. Cells were incubated on cisplatin treatment for 72-hours and analyzed by adding 50µL of CellTiter-Glo 2.0 reagent (Promega) to each well and incubated according to the manufacturer’s instructions. Luminescence was read using a BioTek Cytation 5 Cell Imaging Multi-mode Reader (Agilent). The cisplatin IC_50_ was determined for each cell line by normalizing treatments to the vehicle control and plotting the data in GraphPad Prism 9.

### RNA-sequencing

Total RNA was extracted from cell pellets using a Quick DNA/RNA Miniprep Plus Kit (Zymo Research) per manufacturer’s instructions with the optional on-column DNase digestion and frozen at −80°C until use. RNA quality was assessed using RNA High Sensitivity ScreenTapes and Reagents and analyzed on a TapeStation 4200 instrument (Agilent). The RNA concentration for each sample was quantified using a Qubit^TM^ RNA Broad Range Kit (Thermo Fisher Scientific) according to the manufacturer’s protocol and read on a Qubit^TM^ Flex instrument. cDNA library prep was performed using the KAPA mRNA HyperPrep Kit (Roche) according to manufacturer’s directions with 300ng input and the following modifications: RNA fragmentation at 94°C for 6 minutes, 10 cycles of PCR amplification. Libraries were quantified using a Qubit^TM^ 1x dsDNA High Sensitivity Kit and average fragment size determined using High Sensitivity D1000 ScreenTapes and Reagents and analyzed on a TapeStation 4200 instrument. Libraries were equimolarly pooled to 750pM loading concentration. Sequencing was performed on a NextSeq 1000 P2 100 cycle cartridge according to Illumina protocols. For all RNA-seq experiments, the same library pool was sequenced over multiple runs to achieve the necessary level of sequencing depth.

### RNA-sequencing analysis

BCL to FASTQ conversion was performed on instrument using Illumina’s DRAGEN DRAGEN BCL Convert v3.8.4 for NextSeq 1000/2000. FASTQ files were then used as input into the NextFlow (v22.04.5^20^) pipeline nf-core/rnaseq pipeline (v3.8.1^21^) using genome GRCh38 for alignment and annotation. From pipeline reports of sequencing QC, aligned reads ranged from 25.7 - 40.6 million paired-end reads (mean: 34.4M, standard deviation +/− 3.3M).

Subsequent analysis was performed on the Salmon gene count matrix. Principal component analysis (PCA) was performed with the DESeq2::plotPCA command on Salmon counts transformed using DESeq2::vst to determine if any likely sample swaps existed, as well as to confirm the consensus subtypes inferred from the literature. Data was plotted with ggplot2.

Differential analysis was performed separately for each sensitive/resistant pair of cell lines, using DESeq2::DESeq with model design ∼ Cell line + Replicate. Genes that were differentially expressed with an adjusted p value < 0.05 were designated as differentially up/down regulated genes. Subsequently, gene ontology was performed using clusterProfiler to determine which of the GO biological pathways were overrepresented in the differentially expressed genes (considering up/down regulated genes separately). Finally, Revigo’s rrvgo package was used to cluster and summarize the overrepresented pathways.

### Copy number signature analysis

Copy number signatures were calculated using previously characterized methods and signatures^22^. We applied this to publicly available copy number data for all ovarian cancer cell lines from the Cancer Cell Line Encyclopedia (CCLE) project, as obtained from DepMap. Specifically, ABSOLUTE copy number data (CCLE_ABSOLUTE_combined_20181227) was used, as this is the latest version of copy number data that provides segment means for all chromosome positions, rather than arm-or gene-level values. HS178T and OC316 were removed due to known contamination of these samples with other cell lines. The CCLE data was input into SigProfilerMatrixGeneratorR (v1.2)^23^, which produces a mutational matrix for the set of samples. This matrix was then the input for SigProfilerAssignmentR (v0.0.23)^24^ using the cosmic_fit() function. After this, copy number signatures 1 through 24, as specified by the Catalogue Of Somatic Mutations in Cancer (COSMIC), were grouped based on similar etiology as follows: changes in ploidy, chromothripsis associated amplification, focal loss of heterozygosity, chromosomal loss of heterozygosity, tandem duplication and homologous recombination deficiency, unknown, and profile oversegmentation. Weighted and unweighted copy number signatures were plotted; weighted signatures were generated by summing the unweighted scores and adjusting the fraction of contributions of each score out of 100% of the sum.

### Oncoprints

Mutation and gene copy number data was obtained from cBioPortal using the “CCLE Broad, 2019” dataset. Ovarian cell lines were selected for OncoPrinter plotting. Genes selected were known homologous recombination deficiency genes (from review^25^). For the associated supplemental table, known allele fractions were obtained from DepMap for single nucleotide polymorphisms, as well as amplifications and deletions. Custom tracks were imported into the OncoPrinter tool, including our cisplatin and carboplatin IC_50_ values, fraction copy number altered and scarHRD scores (as described below).

### scarHRD

Homologous recombination deficiency scores were calculated using the scarHRD R package (v0.1.1)^26^. ABSOLUTE copy number data (CCLE_ABSOLUTE_combined_20181227) was obtained from DepMap. scar_score() was run with the following parameters: reference = “grch37”, seqz=FALSE, ploidy = TRUE, chr.in.names = FALSE. The sum of the 3 HRD scores was used as the “scarHRD” score.

### Ploidy and Fraction Copy Number Altered

Ploidy values for cell lines were obtained from DepMap ABSOLUTE analysis from the 20181227 dataset of the CCLE cell lines. Segtab data was used to calculate fraction copy number altered, as follows: sum of the length of regions with copy number alterations that Modal_Total_CN deviated from a value of 2 over the length of all regions measured.

### Generation of Cisplatin resistant OVCAR3 and OVCAR4 cell lines

OVCAR3 and OVCAR4 isogenic platinum-resistant cell lines were generated by treating each cell line with 1uM cisplatin for 4 hours. After treatment, cells were provided with fresh medium and allowed to recover until the cells resumed their typical growth rate for at least two passages (typically about 2 weeks). The same treatment process was then repeated at the same concentration, and after recovery of the second treatment, the cisplatin dose was increased by 1uM and the process repeated. This process was continued over a period of 9 months. Two independent platinum resistant cell lines were generated for both the OVCAR3 and OVCAR4 lines designated ResA or ResB. The cells are not maintained in cisplatin containing medium.

### Flow Cytometry

Cells were harvested using PBS, pH 7.4 + 5mM EDTA to preserve cell surface proteins. Cells were stained (1:50 dilution) with a CD133-APC conjugated antibody (Miltenyi Biotec, cat. 130-113-184) and ALDH activity assessed using an ALDEFLUOR kit (STEMCELL technologies) according to the manufacturer’s protocol. Mouse IgG2b-APC (Miltenyi Biotec, cat. 130-122-932) and single stain controls were performed. Cells were imaged on a CytoFLEX benchtop cytometer (Beckmam Coulter) and data analyzed using FlowJo. A minimum of two independent replicates were performed for all flow cytometry, and the reported ranges reflect the consensus of the experimental values.

### Tumorsphere Culture

Cells were incubated with tumorsphere medium consisting of Dulbecco’s Modified Eagle Medium/F12 (Gibco), B27 supplement (Gibco), Human Recombinant Basic Fibroblast Growth Factor (STEMCELL technologies), Human Epidermal Growth Factor (PEPROTECH), Human Recombinant Insulin (Gibco) and Bovine Serum Albumin (Sigma) in an ultra-low attachment 6-well plate for 1-2 weeks. When tumorspheres were ready to be passaged, the contents of the wells were transferred to a 15mL conical tube using a 5mL serological pipet. The well was rinsed with 1000uL of tumorsphere medium. Cells were incubated for 10-15 minutes to allow them to settle to the bottom of the tube. The supernatant was removed without disturbing the pellets before 0.05% Trypsin-EDTA was added to break the tumorspheres apart. The tumorspheres were incubated for 3 minutes and then pipetted up and down a few times to help break up the tumorspheres. Media containing serum was utilized to inhibit trypsin activity and the cells were centrifuged for 5 minutes at 400rcf at room temperature. Supernatant was removed, fresh tumorsphere media was added, and the cells were plated in an ultra-low attachment 6-well plate.

### Western Blotting

For cisplatin treated samples, cell lines were treated with their respective IC_50_s for 72 hours before being harvested into pellets. Proteins were extracted from cell lines using a 9M urea extraction buffer (9M urea, 4% CHAPS, 0.5% IPG buffer, 50mM DTT) according to the standard protocols. Fifteen µg protein was subsequently loaded per well onto NuPage 4-12% Bis-Tris gels (Invitrogen) and then transferred to PVDF membranes. Fluorescent detection blots were blocked in 5% skim milk in TBST for 1 hour at room temperature and subsequently probed for a housekeeping gene for 1 hour at room temperature. Following incubation, the blots were probed with a primary antibody for the protein of interest overnight at 4°C. The blots were incubated with secondary antibodies (LiCor #926-32211 and #926-68070) at 1:20,000 for 1 hour at room temperature and scanned on Li-Cor CLx. Primary antibodies: CD133 (Abcam ab19898), ALDH1 (BD Biosciences #611195), Actin (Thermo Fisher Scientific #MA5-15739), GAPDH (Cell Signaling Technology #5174S).

### Wound Healing Assays

Cell lines were plated in their respective culture media in a 6-well tissue culture plate and were seeded at a density that would reach 70-80% confluence after 24 hours of incubation. The densities used range from 3×10^5^ to 1×10^6^ cells per well depending on the cell line and the rate of growth. Cells attached to the bottom of the plate for 24 hours before staining with 10uM final working concentration of CellTracker Green CMFDA dye (Invitrogen). After 30 minutes of incubation, a vertical line was manually scratched into the cell layer, using a 1000uL pipette tip. The scratch was imaged on a Cytation 5 with Biospa microscope at 4x magnification at 0hr, 24hr, 48hr, and 72hr. The width of the scratch was measured at each timepoint on the Cytation 5 with Biospa microscope (Agilent).

### Resource Availability

#### Lead Contact

Further information and requests for resources and reagents should be directed to and will be fulfilled by the lead contact, Dr. Jessica D. Lang.

#### Materials Availability

Isogenic cell lines developed in this study are available for use by the community by direct contact with the corresponding author.

#### Data and code Availability

Gene expression data will be made available through GEO at time of acceptance following peer review. Code used for our analysis is deposited at our GitHub repository: https://github.com/jessicalanglab/ovarian_cancer_cisplatin_response_manuscript/.

## Results

### Selection of OvCa cell line panel for analysis

We selected a panel of 36 ovarian cancer cell lines for this study, based on representativeness across a wide variety of ovarian cancer histotypes (**Table 1**). We extensively examined the literature on these cell lines, including the original literature on the tumors the cell lines were generated from, where available, and recent genomic, transcriptomic, and proteomic analyses comparing cell lines and tumors of different histotypes (5–8) to determine the current consensus (**Table 1**). We made the decision not to include some extensively used cell lines, including A2780 and OVCAR5, as these cell lines have been characterized as potentially problematic in recent multi-omic publications. A2780 lacks characteristic TP53 mutations and high copy number alteration observed in HGSOC, the presumed subtype, and based on gene expression, clusters with lung and liver cancer cell lines^16^. Two other studies support the classification of A2780 as Endometrioid Carcinoma^17,19^. However, these studies did not compare A2780 to non-ovarian cancers. OVCAR5 was determined to be a gastrointestinal cancer origin based on gene expression and pathology review^27^. Therefore, to avoid drawing conclusions on cell lines not well representative of patient tumors, we decided to better characterize less used, but more characteristic cell lines.

**Table 1.**
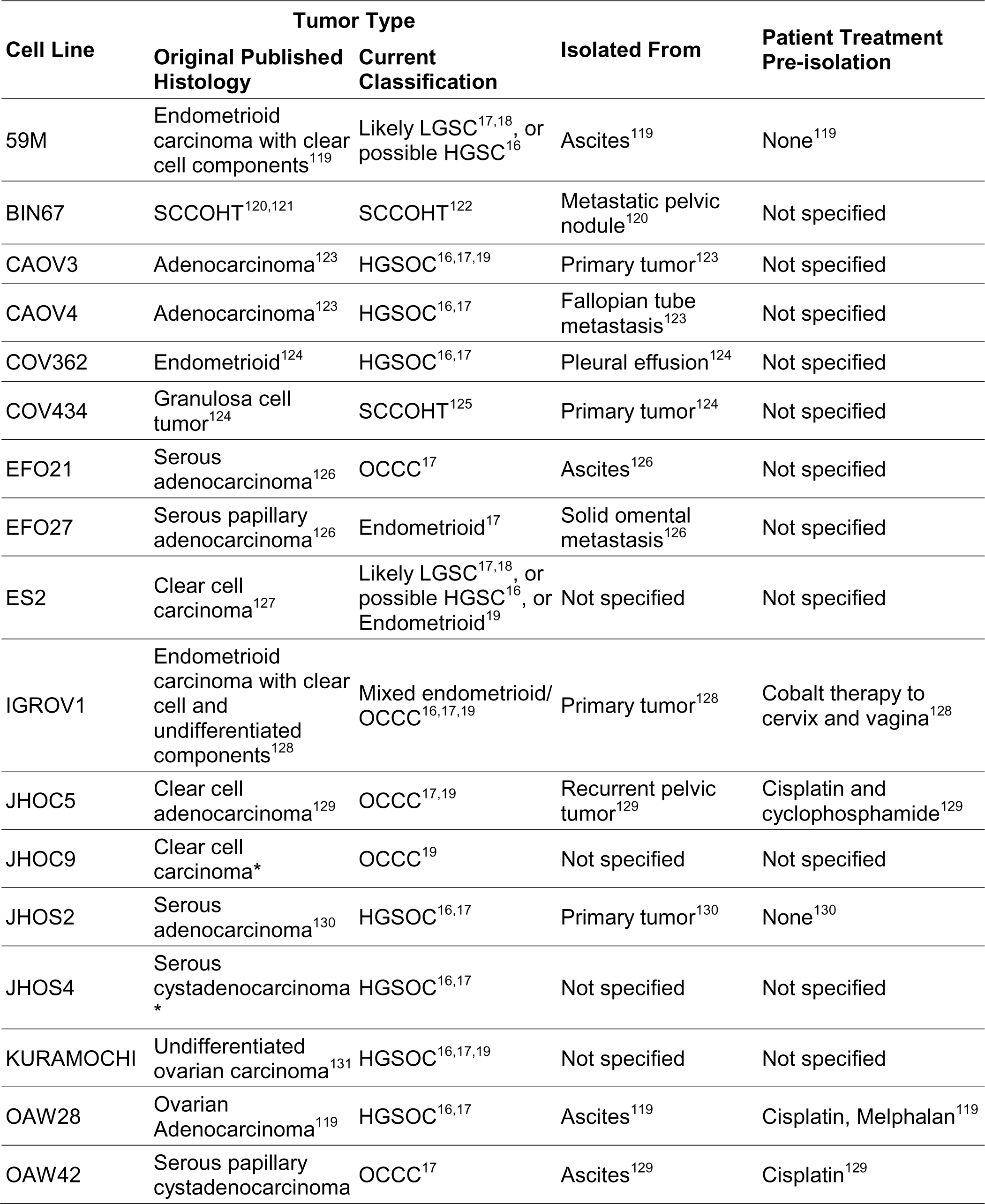
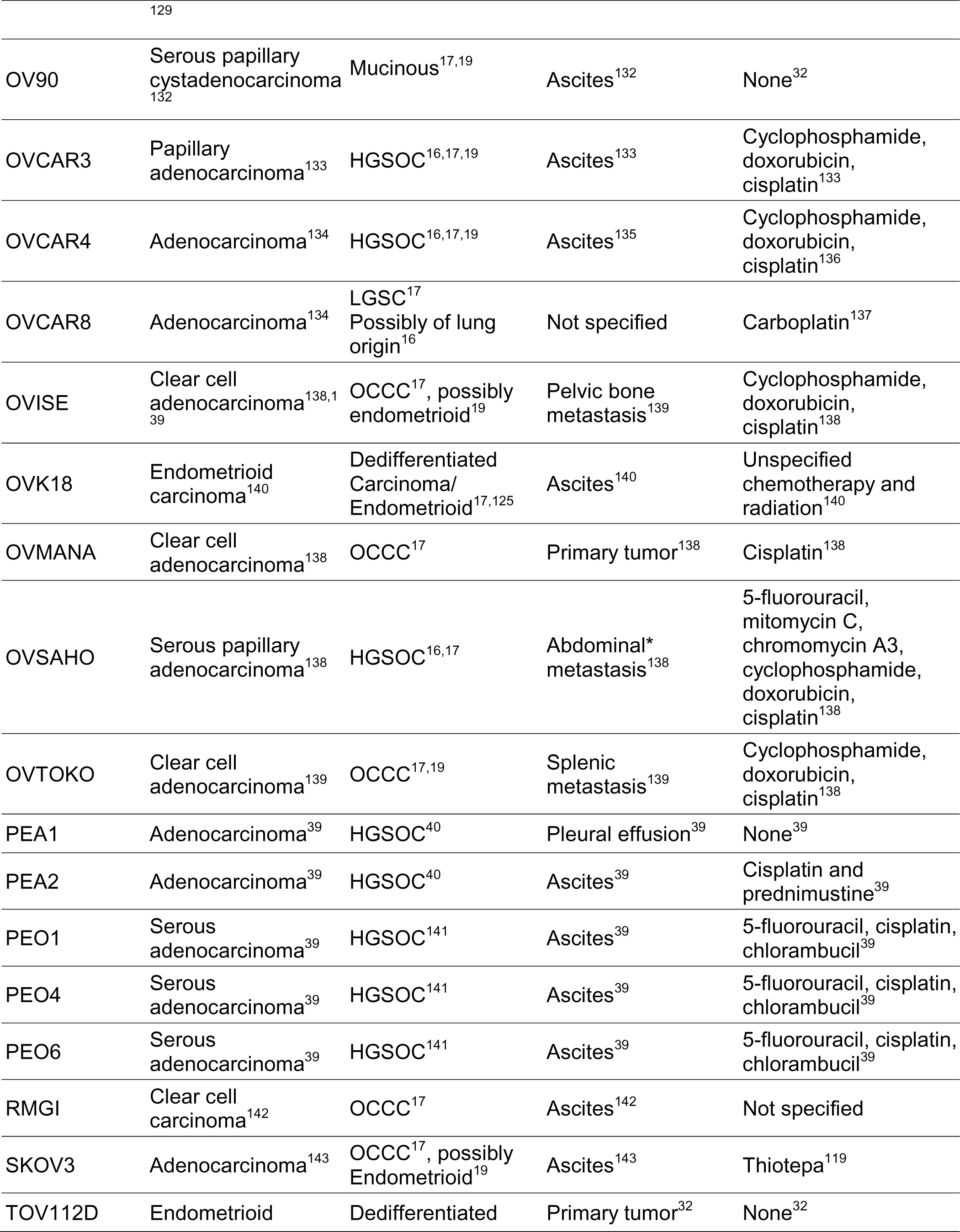
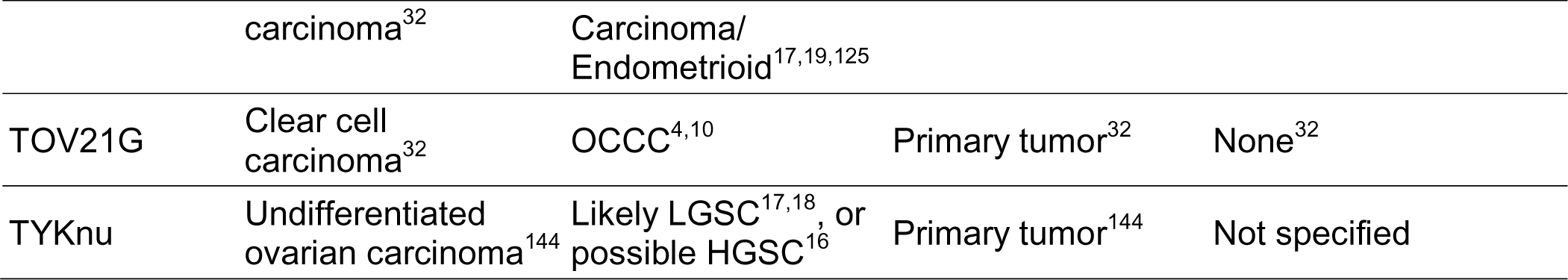
Summary of cell line subtype classification from literature review. *Detail not found in published data but stated in information provided by the cell line supplier.

Where literature was uncertain or disagreed, we also consulted DepMap’s Celligner to compare cell lines to TCGA tumors, which are predominantly HGSOC^28^. Further, we generated RNA-seq data in all 36 cell lines in a single pooled library to allow direct comparison of gene expression with no batch effects. Our final subtype calls from the literature review are supported by the Principal Component Analysis (PCA) clustering based on gene expression (**Figure 1**). The first principal component (PC1) largely discriminates HGSOC and OCCC from other subtypes, and PC2 separates HGSOC from OCCC. Of the 500 genes with the most variance in our dataset used to construct the PCA plot, top ranked features in PC1 included a number of genes biologically relevant to epithelial ovarian cancers. These genes include MUC16, PAX8, and many epithelial cell markers (EPCAM, KRT19, KRT7, KRT8, CDH6, CDH1). The top 20 contributors for each principal component are shown in Figure 1B.

**Figure 1.**
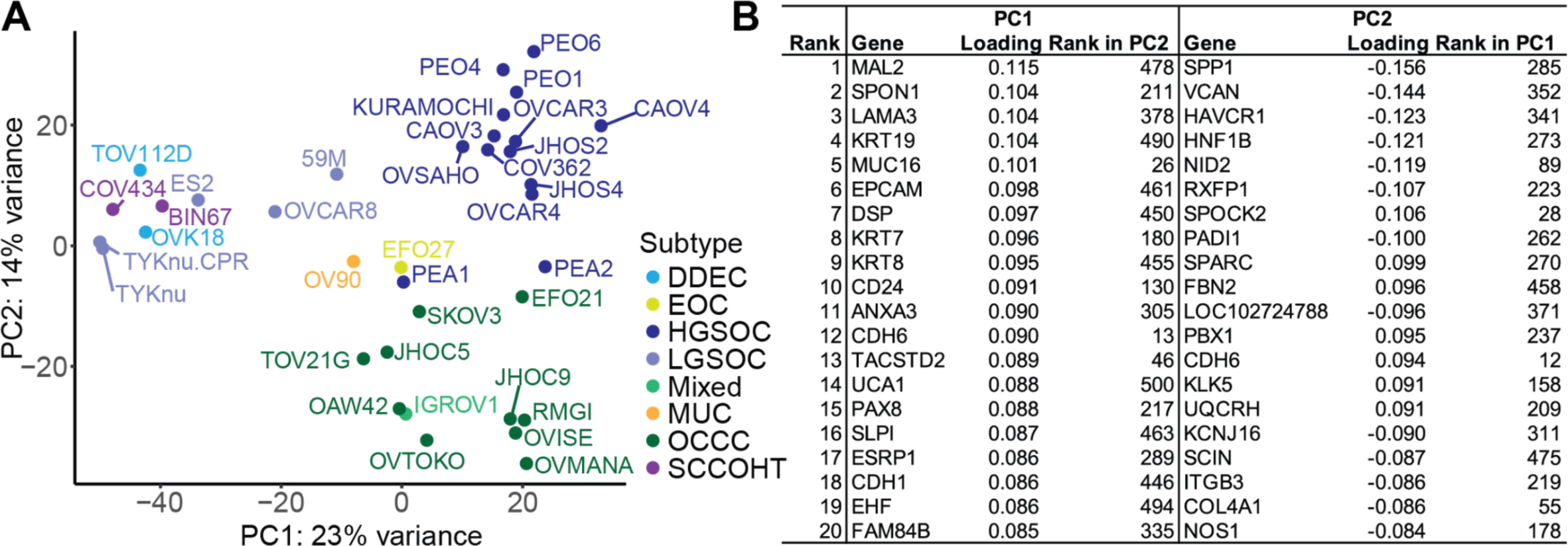
Principal component analysis (PCA) on gene expression across 36 ovarian cancer cell lines included in the study. (A) PCA plot of the first two principal components (PC1 and PC2). Color coding represents the literature-supported subtype classifications. Subtype abbreviations are: DDEC = Dedifferentiated Endometrial Carcinoma, EOC = Endometrioid Carcinoma, HGSOC = High-Grade Serous Ovarian Carcinoma, LGSOC = Low-Grade Serous Ovarian Carcinoma, MUC = Mucinous, OCCC = Ovarian Clear Cell Carcinoma, SCCOHT = Small Cell Carcinoma of the Ovary, Hypercalcemic Type. (B) Top 20 contributing genes to each of the first two PCs. Loadings were ranked by absolute value for each PC, and the corresponding loading rank for each gene in the reciprocal principal component is shown.

### Copy number signature analysis

We performed copy number signature analysis on publicly available genomic copy number data from the Cancer Cell Line Encyclopedia^29^ to examine patterns of copy number variant etiologies, as ovarian cancers, and in particular HGSOCs, have high focal copy number variation^30,31^. From this analysis, a large proportion of the attributed copy number signatures were of unknown origin (CN18-21). The second most frequent etiology was from signatures that indicate focal loss of heterozygosity (LOH; CN9, CN10, and CN12), which occurred most frequently in cell lines presumed to be HGSOC, but also in OCCC cell lines SKOV3, EFO21, and JHOC5, LGSOC cell lines OVCAR8, 59M, and TYKNU, and Mucinous cell lines MCAS and OV90.

Homologous recombination deficiency (HRD) signatures were found in twelve cell lines examined. Of these, seven of them (58%; SNU119, OVCAR4, 59M, KURAMOCHI, OVISE, DOV13, and SNU8) lacked biallelic loss in genes attributed to HRD (**Figure 2** and **Supplemental Table 2**), and occurred in various different subtypes. Scar HRD scores were also applied to the copy number data, and generally overlapped cell lines with HRD called from the copy number signature analysis (**Supplemental Figure 1A** and **Supplemental Table 2)**. CN1 corresponds to diploidy, and CN2 corresponds to tetraploidy; these signatures very closely aligned to reported ploidy of the cell lines (**Supplemental Figure 1B** and **Supplemental Table 2**). HGSOC cell lines had a surprisingly low frequency of tetraploidy-associated signatures (15%), suggesting that copy number signature analysis did not perform well for this subtype, as this contradicts the established etiology of HGSOC. Chromothripsis signatures were infrequently found in ovarian cancer cell lines, and were limited to HGSOC and OCCC cell lines SNU119, EFO21, RMGI, DOV13, and CAOV4. Chromosomal LOH was only found in TOV112D, where it was the only attributed copy number signature. These data are low confidence, as these two cell lines had the lowest total assignments of activities (**Supplemental Fig 1C**). In fact, TOV112D has been described to be hyperdiploid, with 52 chromosomes, including gains in all or parts of chromosomes X, 1, 2, 9, 12, and 15 and 17^32^, and has not previously been described to have chromosomal LOH. However, the reported ploidy score for TOV112D indicates a ploidy of 1 (**Supplemental Figure 1B** and **Supplemental Table 2**).

**Figure 2.**
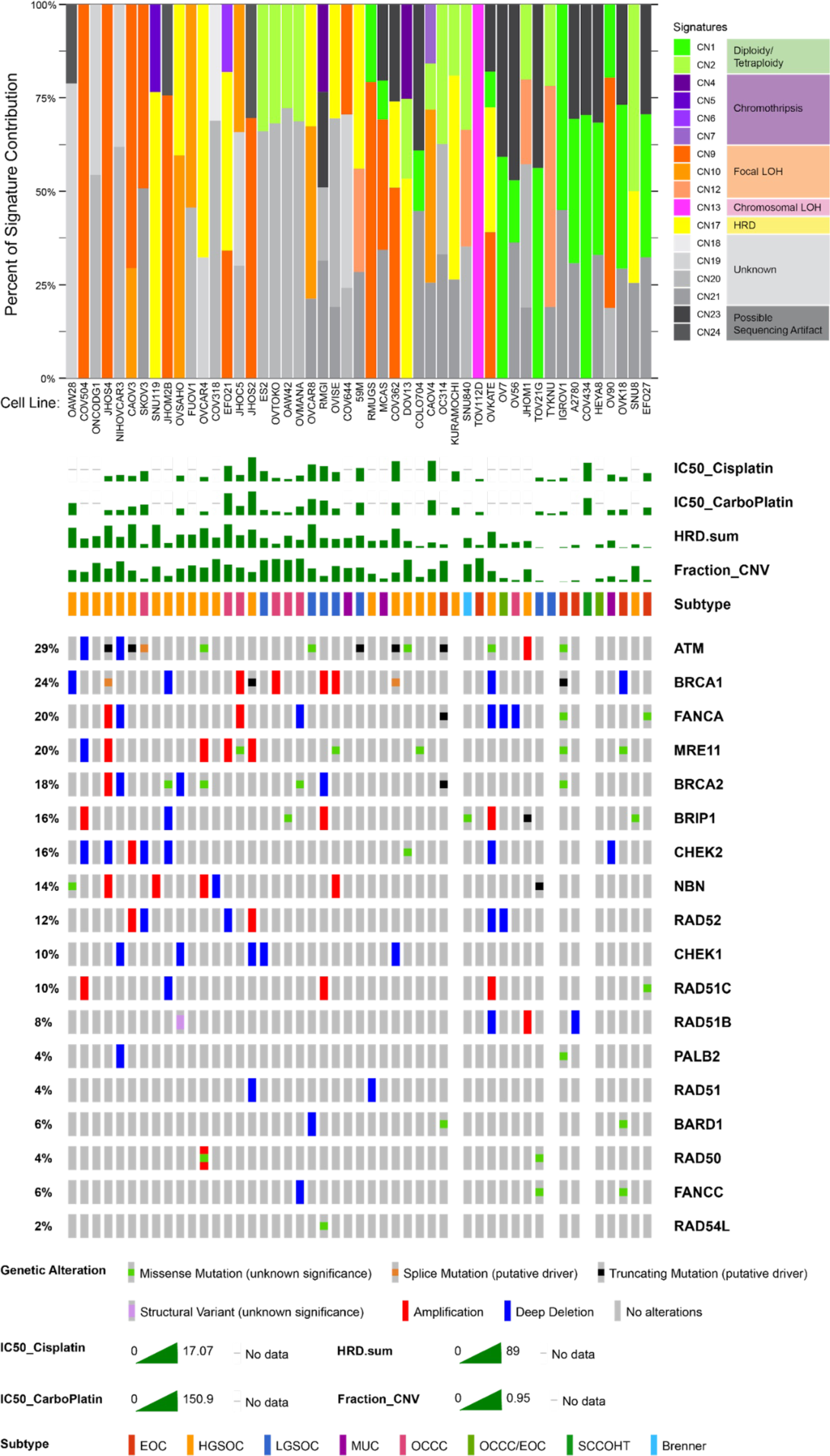
Copy number signature analysis and homologous recombination deficiency (HRD) gene oncoprint for ovarian cancer cell lines in CCLE dataset. The stacked bar plot (top panel) summarizes the weighted contributions of each copy number signature to the overall genomic profile. Copy number signatures are colored by shared etiologies, as described in figure legend to the right. The oncoprint (bottom panel) displays known mutations to HRD genes corresponding to the cell lines in the top panel. IC_50_ values for cisplatin and carboplatin in our own assays are also shown as a bar graph, as well as calculated HRD score (HRD.sum) and fraction genome altered (fraction_CNV). Percentages on the left summarize the fraction of cell lines carrying an alteration to each gene. Subtype is summarized based on our literature review.

### Sensitivity of ovarian cancer cell line panel to platinum-based chemotherapies

For our panel of 36 cell lines, we evaluated the platinum sensitivity of all cell lines by determining the 72-hour IC_50_ values for both cisplatin and carboplatin **(Figure 3A & 3B, Supplemental Table 3A)**. The majority of our cell lines are classified as either HGSOC, LGSOC, or OCCC, and each of those subtypes displayed a broad range of sensitivity to platinum reagents. Between cisplatin and carboplatin, trends in sensitivity generally were similar. For determining relative platinum sensitivity for the purposes of downstream assays, the cells were deemed platinum sensitive within a subtype if their cisplatin 72-hour IC_50_ fell below the SEM, intermediate if it fell within the SEM, and resistant if above the SEM. The range of values for the cell lines determined to be sensitive or intermediate is consistent with clinical C_max_: 58.4-87.4uM for 15-30 minutes of carboplatin infusion at AUC 5-6^33–35^, and 4.1uM and 6.6uM for 1 and 2 hour infusions with cisplatin at 50-80 mg/m^2^ ^36–38^.

**Figure 3.**
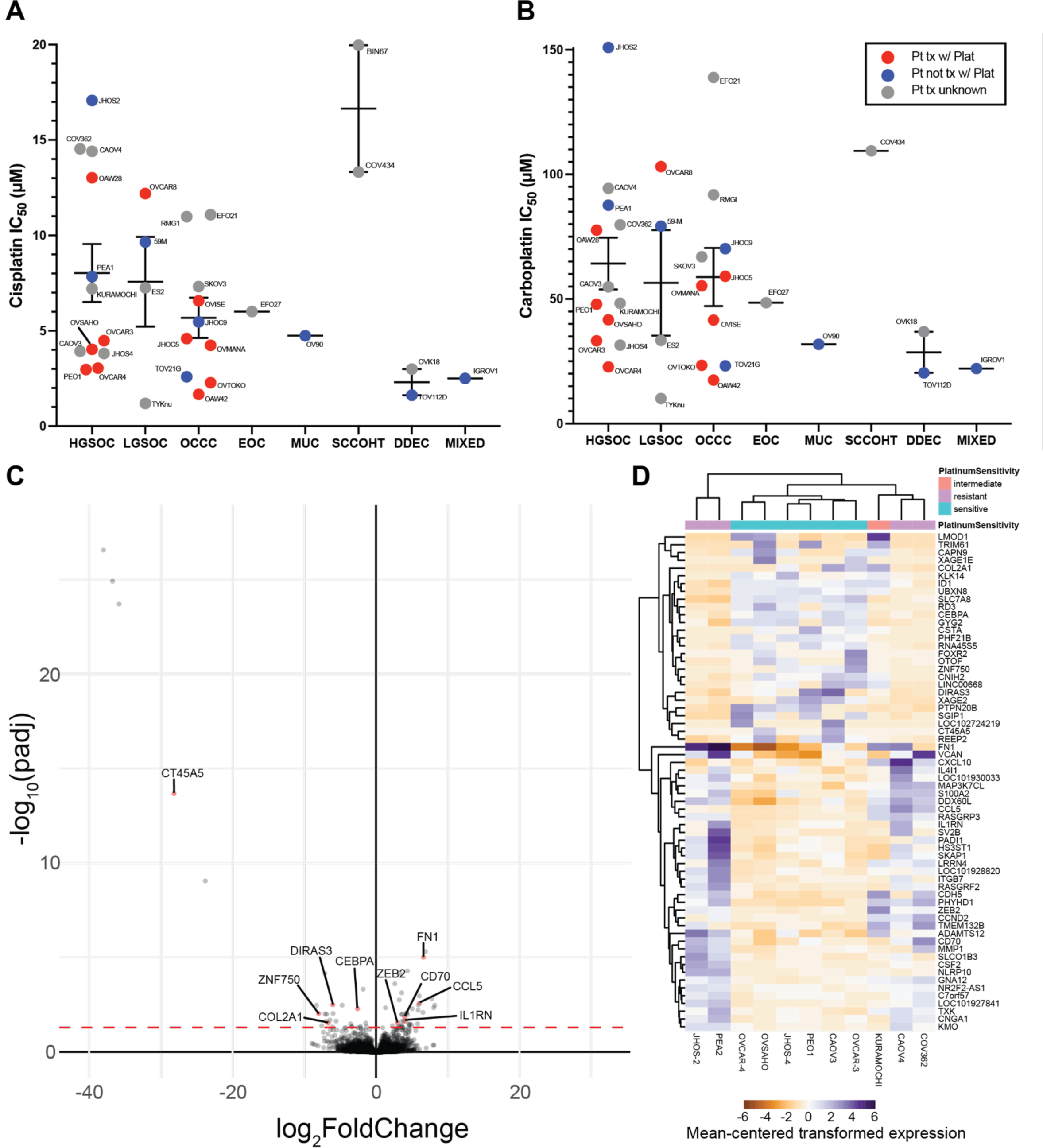
Sensitivity of ovarian cancer cell line panel to platinum-based chemotherapies and resistance-associated gene expression signatures of HGSOC. (A) Cisplatin and (B) carboplatin IC_50_ values for 33 ovarian cancer cell lines of various subtypes. Line and error bars for each subtype show the mean +/− standard error of the mean. (C) Volcano plot and (D) heat map showing differentially expressed genes between HGSOC cell lines based on platinum sensitive, intermediate, and resistant classifications from our data in (A) and (B). Genes with demonstrated roles in platinum-resistance in the literature are labeled in the volcano plot. Values in the heatmap for each gene are centered on the mean of the transformed expression across the eleven cell lines.

### High-grade serous ovarian carcinoma gene expression associated with resistance

Based on the established *in vitro* sensitivity to cisplatin and carboplatin, we used the HGSOC cell lines subsetted into platinum resistant (JHOS-2, PEA2, CAOV4, COV362), intermediate (KURAMOCHI), and sensitive (OVCAR-3, OVCAR-4, OVSAHO, JHOS-4, PEO1, CAOV3) groups to perform differential expression analysis. Only the HGSOC subtype was used for differential expression analysis to avoid finding differentially expressed genes that were related to subtype variations instead of resistance. This resulted in 64 differentially expressed genes, with 37 up-regulated and 27 down-regulated (**Figure 3C, Supplemental Table 3B**). While some genes were consistently up or down in their respective response groups, others were only up or down in a subset of cell lines, resulting in clustering of CAOV4 and COV362 closer to sensitive cell lines (**Figure 3D**).

Of the 56 differentially expressed genes, ten of these had some prior evidence of connection to platinum-resistance. We excluded genes associated with adaptive immune responses, as the conditions under which platinum-resistance was examined in our experiments lacked immune components. The five upregulated genes were CCL5, FN1, CD70, IL1RN, and ZEB2, and the five downregulated genes were CEBPA, DIRAS3, COL2A1, ZNF750, and CT45A5.

### Differential expression of HGSOC isogenic platinum-resistant pairs

To further explore genes related to platinum resistance, we generated isogenic platinum-resistant pairs of the HGSOC cell lines OVCAR3 and OVCAR4 through pulse-treatment of the cells with cisplatin until the IC_50_ value for the cell line increased at least 3-fold. Two independent isogenic pairs were generated for both cell lines (OVCAR3/4 ResA & ResB). In addition, commercially available isogenic platinum sensitive and resistant HGSOC cell lines derived from the same patient were utilized (PEO1/4/6 and PEA1/2)^39^. Collectively, the resistant pairs showed a 1.7-4.8-fold increase in cisplatin resistance (**Figure 4A, Supplemental Figure 2A,** and **Supplemental Table 3A**) and a 1.3-4.5-fold increase in carboplatin resistance (**Figure 4B** and **Supplemental Table 3A**). We performed RNA-seq on the parental sensitive and isogenic resistant pairs in triplicate to determine gene expression changes associated with resistance in a single batch experiment. PCA demonstrates that most differences in gene expression are attributed to unique characteristics of the parental cell lines, rather than from overall shared gene expression influences on platinum resistance (**Supplemental Figure 2B)**. There was not a clear set of genes contributing the most to either dimension (data not shown). Gene ontology analysis revealed a number of resistance-related mechanisms (**Figure 4C)**. Many of the identified pathways were not shared across isogenic pairs, and instead were either only found in a few pairs, or were differentially enriched in different pairs.

**Figure 4.**
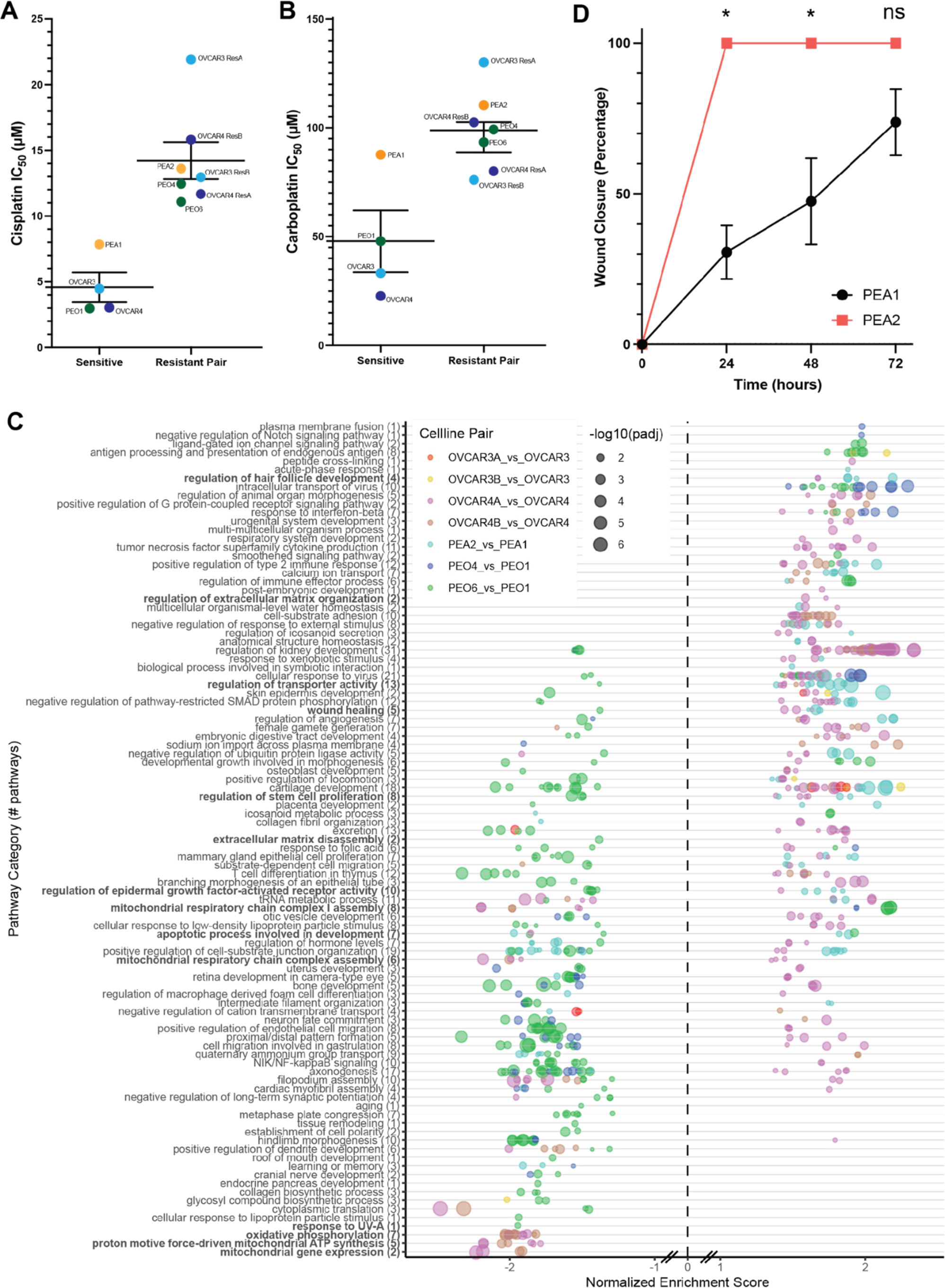
Characterization of isogenic platinum resistant HGSOC cell lines. (A) IC_50_ values for cisplatin and (B) carboplatin comparing the parental platinum sensitive cell line (left) to its isogenic resistant pair(s) (right). Line and error bars for the sensitive cell lines and resistant pairs show mean +/− standard error of the mean. (C) Gene ontology analysis showing overrepresented pathways in the differentially expressed genes found when comparing the sensitive and resistant isogenic pairs individually. Gene ontology results are aggregated by semantic space to shared pathways, where the number of individual gene ontology terms is indicated in the label in parentheses. Each isogenic pair is color-coded independently, as indicated in the legend. (D) Wound healing assay for the PEA1/PEA2 isogenic pair (N = 3 independent replicates). Error bars represent standard deviation. * indicates p-value < 0.05 in paired t-tests and ns indicates non-significant p-value > 0.05.

We layered publicly available annotation database^15^ of genes implicated in platinum resistance on volcano plots in our isogenic pairs to determine whether there was a predominance of key resistance pathways (**Supplemental Figure 3C**). The seven major, non-mutually exclusive, resistance pathways were oncogenic signaling, DNA damage repair, apoptosis inhibition, extracellular matrix modifications, platinum influx/efflux, metabolic changes, and hypoxia responses. At the gene expression level, there is predominance of oncogenic signaling and extracellular matrix modifications as most strongly modified in the direction of contribution to resistance, compared to other mechanism categories. Generally, individual isogenic resistant cell lines from the same parental cell line had more diversity in gene expression contributions to resistance between individual resistant lines than the clinical pair (PEO4/6). This is consistent with clonal ancestry analyses on PEO4 and PEO6 concluding that these were derived from a shared clone^40^.

Of the most dynamically expressed genes known to be associated with platinum resistance, a predominance of Wnt signaling, epithelial-to-mesenchymal transition (EMT), intrinsic inflammatory responses, and platinum influx/efflux were predominant across models. Wnt signaling and EMT related gene expression changes include SOX17, CASP14, KRT5 in OVCAR3 ResB, ANKRD1, MMP10 in OVCAR4 ResA, and EPCAM in PEA2. STAT signaling and innate inflammation-related changes include CD44 in OVCAR3 ResB and OVCAR4 ResB, BIRC3 in OVCAR4 ResA, CXCL12, STAT5B and IL6 in OVCAR4 ResB, and ERBB4, CCL5 in PEO4 and PEO6. Changes to platinum influx and efflux occurred to ABCC3 in OVCAR3 ResB and OVCAR4 ResB, ANXA4, ATP7A in OVCAR4 ResA, and SLC22A3 in PEO4 and PEO6.

Other shared differential gene expression with the highest magnitude of change included GPRC5A (OVCAR3 ResA, PEA2), MAL (OVCAR3 ResA, OVCAR3 ResB, PEA2), S100A9 (OVCAR3 ResB, OVCAR4 ResB, PEA2), SUSD2 (OVCAR3 ResB, PEO4), CLU (PEO4, PEO6), and THBS1 (PEO4, PEO6). 274 genes were differentially expressed in the same direction in five or more isogenic pairs, with 137 up and 137 down. Notable upregulated genes include HDAC6, IER5, IL6, IFIT2, and ITGB8.

Since wound healing was a significant pathway in a few platinum-resistant HGSOC cell lines, we performed scratch assays isogenic platinum-resistant cell lines. Scratch closure was significantly increased for PEA2, OVCAR3 ResA (24h), OVCAR3 ResB (72h) (**Figure 4D** and **Supplemental Figure 4**). The most prominent change was in PEA2, which was consistent with the increase in gene ontology pathway enrichment scores for this cell line (**Figure 4C**). OVCAR4 ResB had significant early delay in scratch closure.

### Stemness markers in isogenic resistant pairs

As cancer stem cell or quiescent states have been implicated in the development of resistance to platinum-based chemotherapy, we examined whether the isogenic platinum-resistant cell lines exhibited more stemness markers compared to their sensitive counterpart. Differential expression analysis of isogenic pairs revealed that PEO4 and PEO6 had increased “regulation of hair follicle development”, PEO6 had decreased “regulation of stem cell proliferation”, and PEA2 and OVCAR4 ResA had increased “regulation of stem cell proliferation” (**Figure 4C**), suggesting involvement of these pathways in cisplatin resistance in these pairs. We examined expression of the two most reliable markers of ovarian cancer stemness phenotype, CD133 and Aldehyde Dehydrogenase (ALDH) by both western blot and flow cytometry in all of our isogenic pairs (**Table 2, Supplemental Figures 5 & 6**). While we anticipated the resistant cell lines having higher expression of these marks, notably, they were nearly always lower in the resistant cell lines compared to their sensitive counterparts. This was also consistent with the ability of each cell line to form tumorspheres. PEO1 had high CD133 expression and ALDH activity, but PEO4 had markedly decreased expression and activity, and could not form tumorspheres. OVCAR3 had higher tumorsphere formation ability than resistant cell line counterparts, which seems to be most correlated with a decrease in CD133 expression for OVCAR3 ResA and a decrease in ALDH activity for OVCAR4 ResB. OVCAR4 resistant clones had partial to complete loss of CD133, but minimal change to ALDH activity, but tumorsphere formation was largely unaffected. PEA2 resistant cells were the only isogenic resistant line to demonstrate increased tumorsphere formation, which correlates with increased ALDH activity in this cell line. Overall, isogenic platinum-resistant cell lines appear to be poor models of stemness as a feature promoting resistance.

**Table 2.** Summary of stemness markers and phenotypes for platinum-resistant isogenic pair cell lines. For western blot, relative expression of CD133 and ALDH1 are shown. For flow cytometry: - less than 2x negative control, + 2x negative control-30% positive cells, ++ 30%-70%, +++ >70%. For tumorspheres: - for no tumorsphere formation, + for no tumorspheres surviving passaging, +++ for tumorspheres forming for at least 2 passages.

| Cell Line | Western Blot |  | Flow Cytometry |  | Tumorspheres |
| --- | --- | --- | --- | --- | --- |
|  | CD133 | ALDH1 | CD133 | ALDEFLUOR |  |
| PEO1 | +++ | - | +++ | +++ | +++ |
| PEO4 | - | - | + | + | - |
| PEA1 | - | - | + | + | + |
| PEA2 | - | - | + | ++ | +++ |
| OVCAR3 | + | +++ | + | + | +++ |
| OVCAR3ResA | - | +++ | - | + | + |
| OVCAR3ResB | + | ++ | + | - | + |
| OVCAR4 | +++ | + | +++ | + | +++ |
| OVCAR4ResA | ++ | + | ++ | + | +++ |
| OVCAR4ResB | + | - | + | + | +++ |

## Discussion

Ovarian cancer cell lines have more recently come under scrutiny for the suitability of representation of commonly used models for the current subtype classifications. In this study, we contribute to the growing body of literature on reclassification using our gene expression data from a single-batch assay of 36 cell lines. Our RNA-seq data recapitulates the most recent reclassifications of cell lines^16–19^. Genes that contributed to the most variation between subtypes included known biomarkers and master regulators of ovarian cancer subtypes (MUC16 and PAX8) and many keratin and cadherin genes (KRT19, KRT7, KRT8, CDH6, and CDH1). We further examined copy number signatures, as ovarian cancer subtypes differ substantially in their chromosomal instability and aberrations. On one end of the spectrum, HGSOCs have exceptionally high copy number alteration^30,31^, whereas SCCOHT are genomically quiescent at both the copy number and mutation standpoint^41^. Copy number signatures have been previously applied to TCGA and other tumor data, but have not yet been reported for cell lines. Ovarian cell line copy number signature analysis identified heterogeneous etiologies even within a subtype. Yet, focal copy number signatures were common features in HGSOC cell lines, which aligns with current understanding that subsequent to TP53 loss in HGSOC, early genomic instability results in genome doubling and subsequent focal copy number alterations. However, not all HGSOC cell lines displayed signatures or ploidy estimates consistent with genome doubling. Outliers to the genome doubling hypothesis include JHOS4, COV504, CAOV3, RMUGS, COV362, JHOS2, OVKATE, JHOM2B, and COLO704, which appear to be near-diploid by these estimates. Further, the HRD-related copy number signature is generally associated with the computed scarHRD score, but some discrepant cell lines were found between these predictors. CAOV3, JHOS4, and OAW28 had high scarHRD scores, but no HRD copy number signature, whereas OVISE, DOV13, and SNU8 had low scarHRD scores, but presence of HRD copy number signatures. Notably in these discrepant cell lines, the cell lines with high scarHRD had deleterious mutations to BRCA genes (either deletions or truncating mutations), but those with presence of only the HRD copy number signature were wildtype for BRCA genes or carried unknown missense mutations or amplifications.

Ovarian cancer cell lines have long been used to study therapeutic response, yet a comprehensive examination of platinum-response in equivalent experimental setup has not been robustly reported in this many cell line models. To our knowledge, the most similar study examined 39 cell lines for three platinum compounds, but this study only reported visualization of relative sensitivity in a heatmap format, and did not contextualize sensitivity to clinically achievable doses^42^. We robustly examined response to both cisplatin and carboplatin using the same method across 36 different ovarian cancer cell lines that were examined by RNA-seq analysis. IC_50_s generally diverged into values that were either above or below the established C_max_ of cisplatin and carboplatin. Thus we are confident that there is a clinically translatable resistance level in the models we have examined. Interestingly, cell lines of certain subtypes displayed variable platinum sensitivity. HGSOC and LGSOC displayed a wide range of sensitivity, with cell lines spanning platinum doses above and below the C_max_. EOC, OCCC, Mixed, and DDEC cell lines were largely sensitive to platinum. SCCOHT cell lines were exclusively resistant to platinum, which is consistent with the clinical observations that high-dose chemotherapy is most effective for these patients^43^.

Within HGSOC cell lines, of which we had twelve that clustered well together, we performed differential expression analysis and found only a select set of genes previously implicated in platinum resistance. The five upregulated genes were CCL5, FN1, CD70, IL1RN, and ZEB2. While CCL5 is generally a chemoattractant to recruit immune cells, it has been implicated in ovarian cancer resistance through activation of pathways in cells directly, either through local secretion that causes STAT2 and PI3K/Akt activation^44^ or through activation subsequent to BRCA1 inactivation which can lead to STING activation to promote inflammation^45^. FN1 is a fibronectin glycoprotein that has been implicated in inducing cisplatin resistance and correlating with patient outcome in ovarian and other cancers^46–49^. CD70 is a TNF-family cytokine that is typically found on activated T and B cells, but is known to be over-expressed in platinum-resistant A2780 cells^50^ and is associated with poor survival and chemoresistance^51^. IL1RN is an interleukin 1 family member that inhibits IL-1. Lower abundance of IL1RN in ascites is associated with improved overall survival outcomes^52^. ZEB2, a transcriptional repressor, has been associated with epithelial to mesenchymal transition in ovarian cancer^53^ and is re-expressed in models of chemoresistant ovarian cancer^54^.

The five downregulated genes were CEBPA, DIRAS3, COL2A1, ZNF750, and CT45A5. Low expression of the CEBPA transcription factor is associated with poorer overall survival and growth and progression phenotypes in ovarian cancer cell lines, and has been shown to repress WNT signaling^55^. DIRAS3 is a member of the Ras super-family that is an imprinted tumor suppressor gene. Its loss, which occurs in more than half of ovarian cancers, has been associated with reduced autophagy and cisplatin resistance^56,57^. COL2A1, a component of type II collagen, has been shown to positively correlate with higher sensitivity to platinum in HGSOC^58^. ZNF750 is a transcription factor normally associated with epidermis differentiation whose expression is positively correlated with increased sensitivity to chemoradiotherapy in esophageal squamous cell carcinoma^59^. While it has not been implicated in ovarian cancer, it is consistent with a loss of epithelial-mesenchymal transition (EMT) phenotype that is associated with chemoresistance. CT45A5 is one of a cluster of homologous genes that encode the CT45 cancer/testis antigen. CT45 has been implicated in inhibiting DNA damage repair and is more highly expressed in platinum-sensitive tumors^59^.

In addition, we used seven total platinum-resistant isogenic pairs, including three well established isogenic pairs from clinical platinum resistance^39^, and four novel resistant cell lines derived from pulse-treatment of platinum-sensitive OVCAR3 and OVCAR4 cells to cisplatin. The choice of pulse-treatment was based on previous work that demonstrates that this most closely recapitulates selection pressure observed in patients^60^. Again, we found a predominance of innate immunity/STAT activation, EMT, and platinum influx/efflux regulators in the differentially expressed genes between resistant and sensitive cell lines. Wnt signaling and EMT related gene expression changes were predominantly found in OVCAR3 ResB, OVCAR4 ResA and PEA2. SOX17, CASP14, KRT5, ANKRD1, MMP10, EPCAM. SOX17, which is decreased in some resistant lines, is notable because it has been proposed to be a master regulator in ovarian cancer^61^ and inhibits WNT3A-stimulated transcription through degradation of beta-catenin^62^. Downregulation of SOX17 also allows higher expression of DNA damage repair genes^63^, and its expression is generally associated with increased cisplatin sensitivity^63–65^ though this has not been described for ovarian cancers. CASP14 and KRT5 are associated with keratinization, and have been implicated in resistance to lung and ovarian cancers^66–68^. ANKRD1, a transcription factor target of TGF-beta and Wnt signaling is necessary for resistance to platinum, and its expression is associated with poorer outcomes in ovarian cancer^69^. Endoplasmic reticulum stress has also been posited to play a role in ANKRD1-related platinum resistance^70^. MMP10, a matrix metalloproteinase that degrades gelatins and fibronectin is over-expressed in platinum-resistant A2780 cells^71^ and is also thought to activate canonical Wnt signaling through preventing Wnt5A-mediated noncanonical signaling^72^. EPCAM, an epithelial adhesion protein, has an intracellular domain shown to induce Wnt signaling when cleaved. Expression of EPCAM is increased in platinum-resistance and associated with EMT^73^, and high expression is associated with poorer prognosis^74^, however this is not supported by TCGA data^75^.

STAT signaling and innate inflammation-related changes were mostly found in OVCAR3 ResB, OVCAR4 ResA, OVCAR4 ResB, PEO4, and PEO6. CD44, IL6, BIRC3, CXCL12, STAT5B, ERBB4, CCL5 were notable STAT-related gene expression changes. CD44, a receptor for hyaluronic acid, is a cancer stem cell marker frequently associated with chemoresistance that can activate STAT3. Autocrine signaling of the interleukin IL6 has been associated with platinum resistance in many cancers^76–80^, including ovarian cancer^81^. IL6 upregulates HIF-1alpha through STAT3 activation^82^. BIRC3, a E3 ubiquitin-protein ligase that regulates NFkB activation and anti-apoptotic caspase regulation can be induced by IL-6 treatment in ovarian cancer cells^83^. Over-expression of BIRC3 is observed in platinum-resistant A2780 cells^84^ and in response to RUNX3 OE in A2780, which occurs in resistant derivatives^85^. CXCL12, also known as Stromal cell-derived factor 1 is a chemokine that activates CXCR4 and increases intracellular calcium and chemotaxis. CXCL12 treatment of SKOV3 makes them more resistant to cisplatin^86^ and induces Wnt/beta-catenin pathways and EMT^87^. In lung cancer, CXCL12 activates JAK2/STAT3 signaling to mediate resistance to cisplatin^88^. STAT5B, a transcription factor effector of inflammatory signaling, can be activated downstream of ERBB4 and IL-11 to induce resistance to platinum^89,90^ and can be targeted with the STAT5 inhibitor dasatinib to restore sensitivity to platinum^91^. ERBB4, an epithelial growth factor receptor family member, has also been shown to activate Ras/MEK/ERK, PI3K/AKT, STAT3 and STAT5, with high expression associated with poor survival in ovarian cancer^92^. Chemokine CCL5 has been demonstrated to desensitize ovarian cancer cells to cisplatin through expansion of cancer stem cells^93^, and promotes resistance to cisplatin through STAT3 and PI3K/AKT activation^44^.

Changes to platinum influx and efflux were frequent in OVCAR3 ResB, OVCAR4 ResA, OVCAR4 ResB, PEO4, and PEO6. ABCC3, ANXA4, ATP7A, and SLC22A3 were particularly notable examples of platinum influx/efflux changes. ABCC3 is a well known multidrug transporter channel involved in the efflux of many drugs out of cells. ANXA4, or Annexin A4, promotes membrane fusion and exocytosis and is involved in platinum resistance in OCCC^94,95^. Notably, we observe ANXA4 increased in HGSOC in this study. ATP7A is a copper-transporting ATPase 2 that exports copper out of cells and has also been shown to mediate resistance to platinum agents^96,97^. Cation transporter SLC22A3 is involved in influx of cisplatin into cell, but may not transport carboplatin16914559^98^. Decreased SLC22A3 expression is associated with poorer prognosis^99,100^.

Other genes with shared differential gene expression across multiple models with high or frequently shared differential expression were GPRC5A, MAL, S100A9, SUSD2, HDAC6, IER5, IFIT2, and ITGB8. GPRC5A is a marker of poor outcome in HGSOC^101^. MAL, which is involved in vesicular trafficking and lipid raft organization, is generally associated with resistance and poor outcomes^102,103^, and MAL promoter hypomethylation is associated with increased expression of MAL in resistant tumors^104^. S100A9 is more highly expressed in resistant tumors^105–107^, and its increase is evident following acute treatment with neoadjuvant chemotherapy^108^. Sushi-domain containing membrane protein receptor SUSD2’s expression promotes chemotherapy resistance^109^. HDAC6, which was upregulated in all isogenic pairs except PEA2 and OVCAR3 ResB is a histone deacetylase that interacts with DNA damage repair factors in HGSOC^110^. High HDAC6 is associated with poor prognosis and chemoresistance in patients with HGSOC^111^. IER5, whose expression was up in all but the OVCAR4 ResA and ResB lines, is associated with poor disease-free intervals and has been shown to be induced by platinum^112^. IFIT2, upregulated in all but PEA2 and OVCAR3 ResB, has been associated with chemoresistance previously in HGSOC^113^. Its expression is induced by interferon signaling. ITGB8, an integrin beta subunit, was upregulated in all but OVCAR3 ResA and ResB and has been shown to be regulated by noncoding RNAs to mediate stemness and therapy response^114,115^.

While we anticipated seeing that resistant cells would have higher positivity for markers of stemness, this was generally not the case, indicating that stemness phenotypes must be a transient state through which resistant cells pass, not a selected trait. This is consistent with the persister cell hypothesis^116^. Notably, as described above, despite the lack of clear canonical markers of stemness phenotype and function, we do see some gene expression changes to EMT and stemness genes that may contribute to resistance. Our overall conclusion from the stemness assays is that isogenic platinum-resistant cell lines are not a good model of the transitory state of platinum resistance development, but represent the states that likely exist in a steady-state resistant population.

We note some limitations of the design of our study. It is known from other studies that immune responses and stromal interactions are quite important for overall response of tumors to therapy, and we recognize that these were inadequately represented by our study. While we made attempts to compare the cell lines representing diverse subtypes of ovarian cancer to clinical tumor RNA-seq data, a comprehensive dataset including a representative set of each of these tumors was not publicly available. Inspired by CASCAM^117^, we attempted to batch correct several subtype-specific datasets and integrate with our own data using and ComBat^118^, but we found that we were unable to resolve the clustering by dataset without sacrificing clustering by ovarian cancer subtype. Since most studies are predominated by a single subtype, batch correction not only normalized experiment-to-experiment variability in datasets, but also the intersubtype variability in expression (i.e. the gene expression that contributes to subtype identity). Future studies with RNA-seq data evenly representing different ovarian cancer subtypes would aid in building a foundation upon which we could better assess cell lines against patient tumor subtypes.

Our data suggest multi-mechanism contributions to resistance even within a single resistant cell line that continue to be a challenge to tease apart. Future work will quantitatively determine individual mechanisms to resistance. Yet, the robustness of the cell lines chosen and the number of isogenic pairs examined is a strength compared to similar studies on gene expression changes with resistance in ovarian cancer cell models, which have mostly relied on A2780 and SKOV3 derivatives. Further, applying the differential gene expression changes to a clinical transcriptomic dataset with well-annotated platinum-free intervals (PFI) would greatly improve the biomarker implications of our findings, but even TCGA lacks robust numbers of tumors with both RNA-seq and PFI information available (on the order of 40 tumors). In the future, we plan to build on our gene expression networks through the integration of shared master regulators and epigenetic organization to develop a more integrated picture of upstream targets with pleiotropic effects that converge to promote resistance.

## Acknowledgements

Research reported in this publication was supported by the National Cancer Institute of the National Institutes of Health under award number R00CA234391 to JDL.

## Author Contributions

Conceptualization, K.M.A., J.D.L; Data Curation, R.M.; Formal Analysis, J.W., R.M., Z.J., S.G.; Funding Acquisition, J.D.L,; Investigation, K.M.A., J.R.W., J.W., S.O., R.M.; Methodology, K.M.A., J.R.W., J.W., S.O.; Project Administration, J.D.L.; Supervision, J.D.L.; Visualization, K.M.A., J.W., R.M., S.G., J.D.L.; Writing-Original Draft, K.M.A., J.D.L., Writing-Review & Editing, K.M.A., J.R.W., J.W., J.D.L.

## Supplemental Table and Figure Legends

**Supplemental Table 1.** Cell line sources, culture conditions, and assay cell densities for each cell line used in our study.

**Supplemental Table 2.** Homologous recombination deficiency (HRD) scores and ploidy estimates for ovarian cancer cell lines. In the HRD tab, the scarHRD HRD score is reported as HRD.sum. Presence of copy number signatures related to HRD are reported as a value of 1 in the “HRD CN signature”. Mutations to common HRD-related genes are also specified by gene, where numeric values indicate the known allelic fraction. In the ploidy tab, ploidy indicates the value from CCLE data. Presence of copy number signatures related to genome doubling and diploidy/tetraploidy are shown as a value of 1. Fraction genome altered is also reported based on CCLE data.

**Supplemental Table 3.** (A) Cisplatin and Carboplatin IC_50_ values for each cell line used in the study, including isogenic platinum resistant cell lines created in our laboratory. All values are from a 72 hour assay. (B) Differential gene expression analysis of HGSOC cell lines, comparing resistant cell lines to sensitive cell lines, such that a positive log2 fold change indicates higher expression in resistant cell lines. (C) Differential gene expression analysis of isogenic platinum-resistant pairs, such that a positive log2 fold change indicates higher expression in the resistant cell line compared to their sensitive parental counterpart.

**Supplemental Figure 1.** Association of between homologous recombination deficiency (HRD) scores and ploidy estimates for ovarian cancer cell lines. (A) ScarHRD scores are plotted for each ovarian cancer cell line in CCLE, then labeled blue if the cell line also had a homologous recombination-related copy number signature etiology associated with it. (B) Ploidy estimates from CCLE for ovarian cancer cell lines adjusted such that 0 indicates diploidy are plotted. Bars are colored light green for presence of tetraploidy copy-number signatures (CN2, CN10, CN12) and dark green for presence of diploidy copy-number signatures (CN1, CN9). Unweighted copy number signature contributions for each cell line.

**Supplemental Figure 2.** Isogenic platinum-resistant pair drug response and gene expression relatedness. (A) Cisplatin drug dose response curves. Error bars represent standard error of the mean across 3-4 independent replicate experiments, each with 3 technical replicates. (B) Principal component analysis (PCA) based on RNA-seq for each isogenic cell line pair. Individual replicates are shown as individual points, with shape corresponding to platinum-response status (sensitive vs. resistant) and shades of a color corresponding to individual lines in a related isogenic family.

**Supplemental Figure 3.** Volcano plots showing differentially expressed genes in the platinum sensitive/resistant isogenic cell line pairs divided into categories related to platinum resistance. Based on previous literature review of genes associated with platinum-response^15^, genes were categorized based on seven main, non-mutually exclusive mechanisms of resistance, which are plotted individually. Point size indicates the strength of evidence for contribution to platinum resistance, and points are colored green if directionality of change with resistance is positively correlated with the log2-fold change observed in our data, and red if directionality of change was negatively correlated (opposite direction) as the log2-fold change observed in our data.

**Supplemental Figure 4.** (A) Representative images of a scratch wound healing assay for the PEA1/2 isogenic cell line pair. Red lines indicate the manually annotated scratch boundary, where it could be found. (B) Scratch wound healing assay data for the OVCAR3 (N = 4 independent replicates), OVCAR4 (N = 4), and PEO1/4 (N = 3) isogenic cell line pairs. Error bars represent standard deviation. * indicates p-value < 0.05 in paired t-tests and ns indicates non-significant p-value > 0.05.

**Supplemental Figure 5.** (A) Representative ALDH1A1 western blot images of all isogenic cell line pairs along with platinum resistant cell lines JHOS2 and COV362. GAPDH serves as a loading control. (B) CD133 and ALDH1A1 western blot images of all isogenic cell line pairs with or without cisplatin treatment at each respective cell line’s IC_50_ for 72 hours. GAPDH serves as a loading control.

**Supplemental Figure 6.** Representative flow cytometry data for CD133 and ALDEFLUOR (ALDH activity) for each isogenic cell line pair. DEAB is an inhibitor of ALDH enzymes that serves as a baseline negative control in the ALDEFLUOR assay. Negative control for CD133 stain is isotype control. Plot shown were first gated on live cell population.

