## Supplemental figures for "Cell-intrinsic platinum response and associated genetic and gene expression signatures in ovarian cancer cell lines and isogenic models"

### Supplemental Figure 1

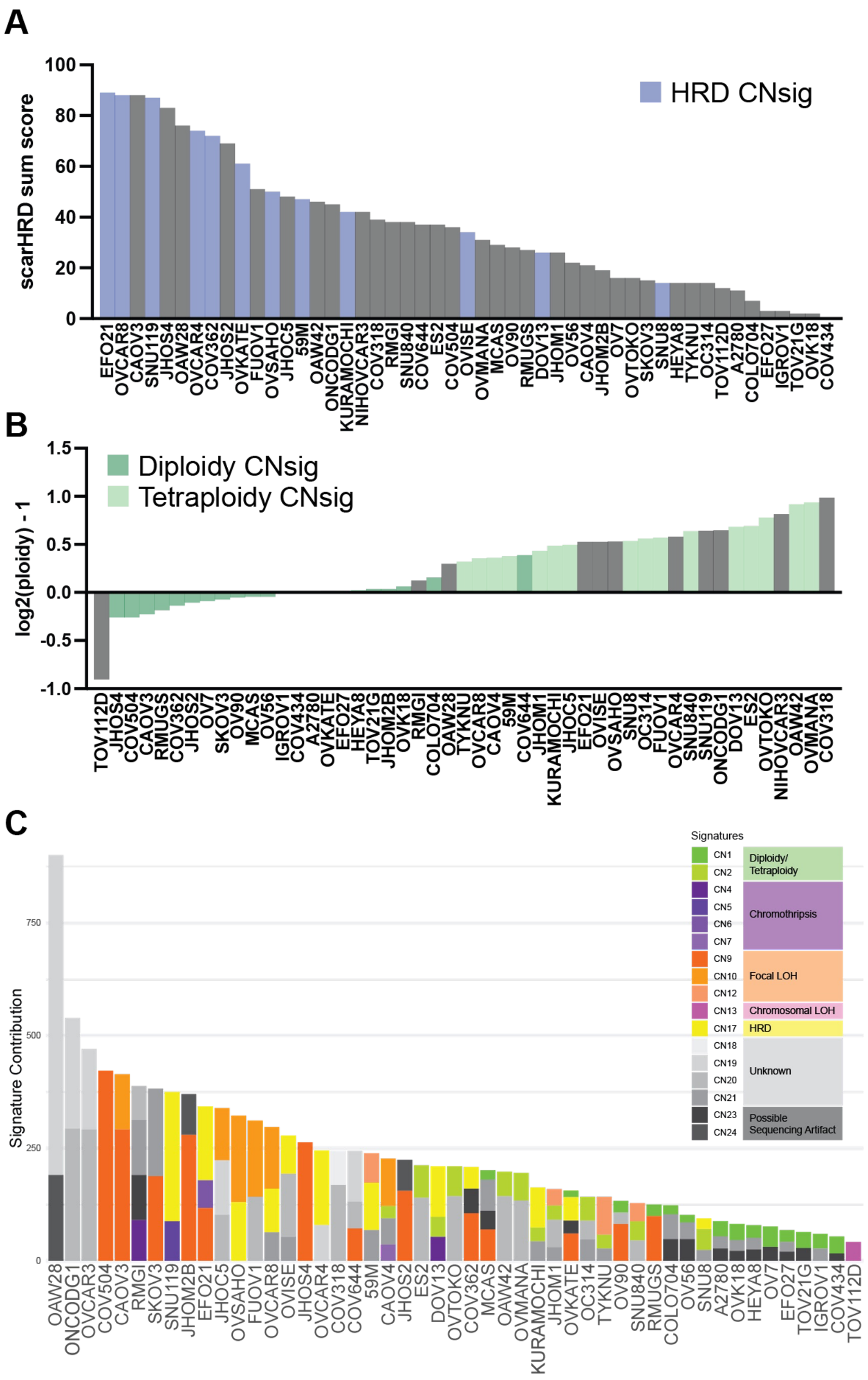

### Supplemental Figure 2

A

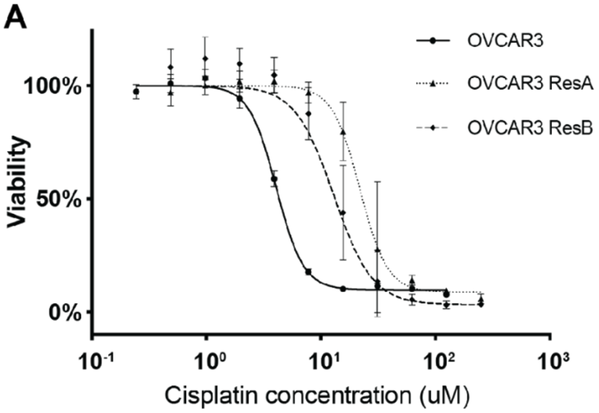


| | OVCAR3 | OVCAR3 ResA | OVCAR3 ResB |
|---|---|---|---|
| IC50 | 4.081 | 21.91 | 12.95 |

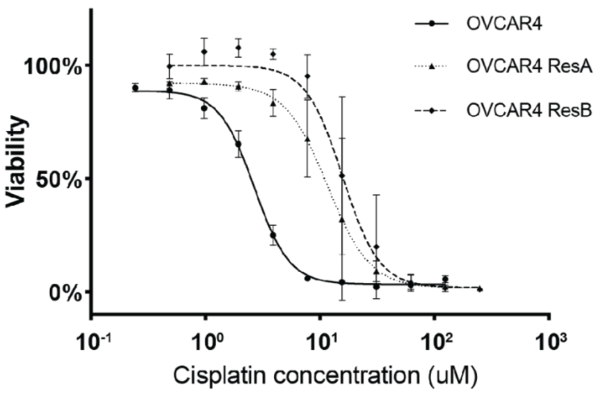


| | OVCAR4 | OVCAR4 ResA | OVCAR4 ResB |
|---|---|---|---|
| IC50 | 2.682 | 11.68 | 15.81 |

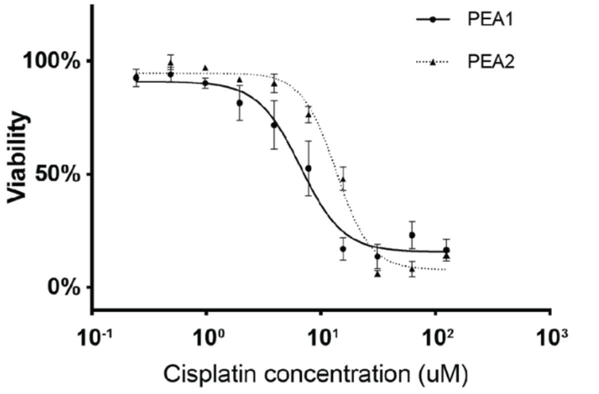


| | PEA1 | PEA2 |
|---|---|---|
| IC50 | 6.640 | 13.62 |

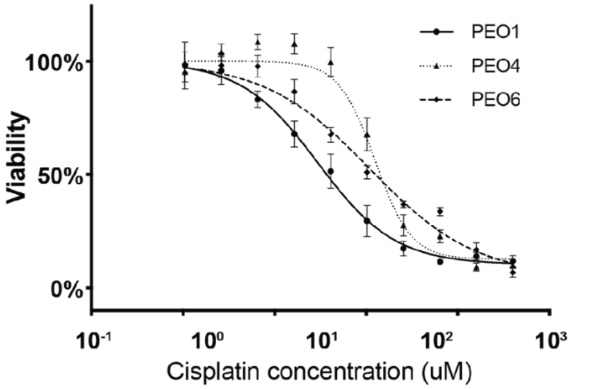


| | PEO1 | PEO4 | PEO6 |
|---|---|---|---|
| IC50 | 2.972 | 12.45 | 11.07 |

B

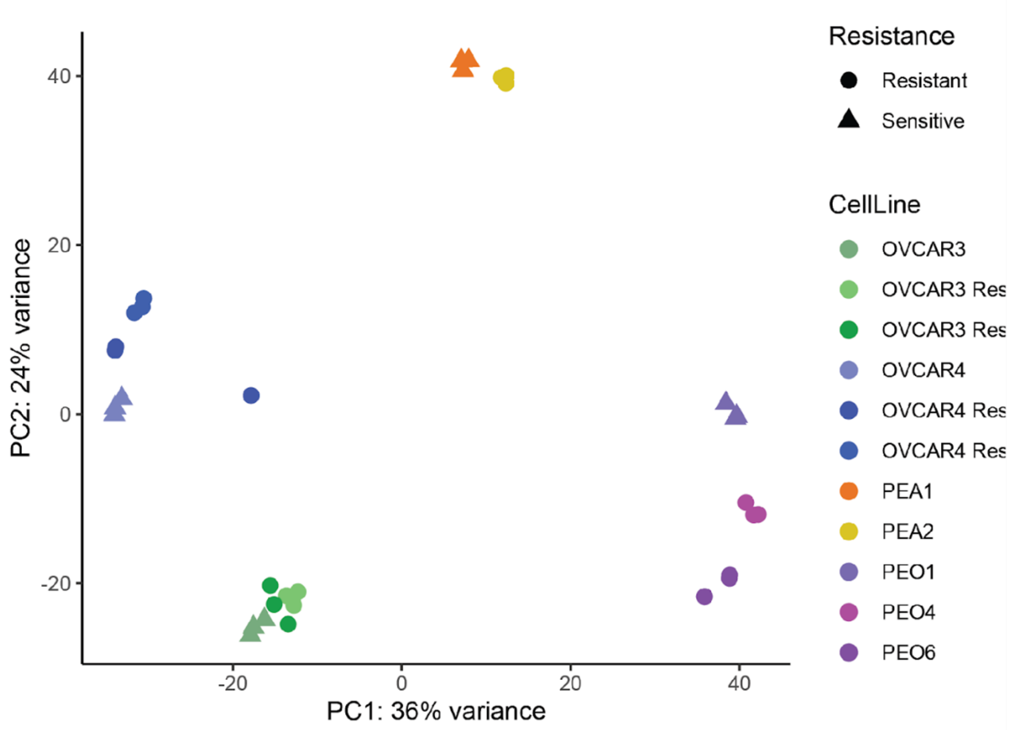

### Supplemental Figure 3 – OVCAR3 ResA (1 of 2)

Enhanced repair and tolerance of platinum induced DNA damage and blockage of cell cycle inhibition

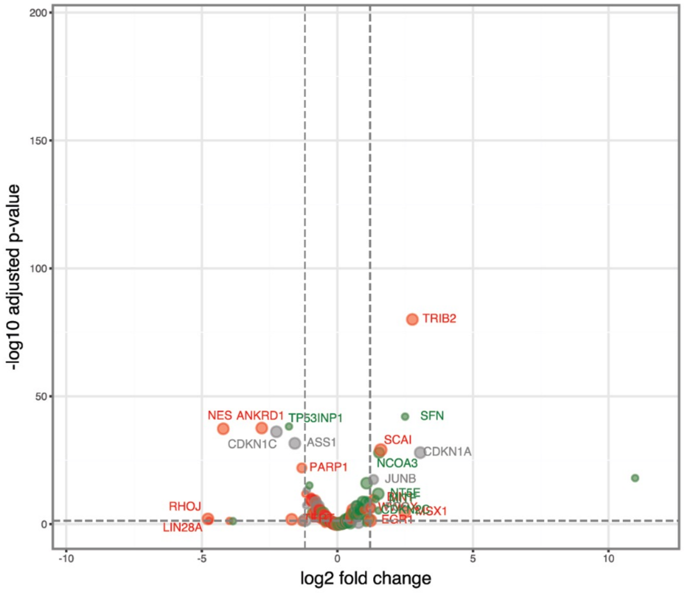


Extracellular mechanisms that alter the extracellular matrix (ECM) and enhance tumor-promoting inflammation

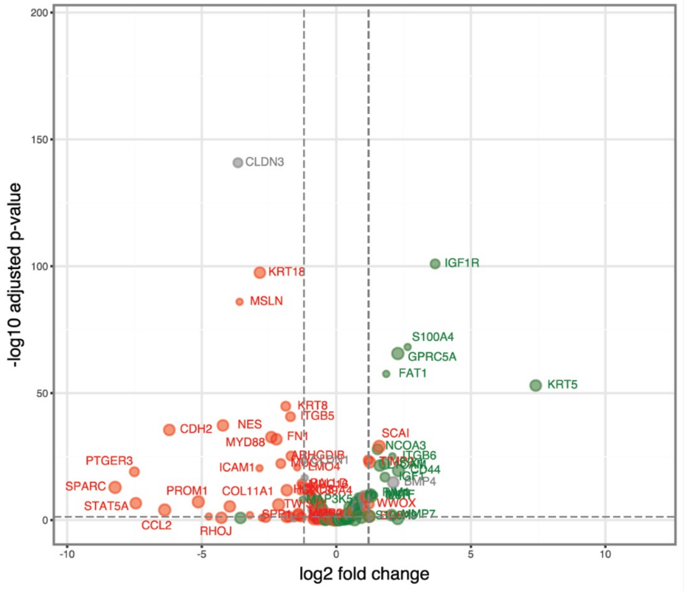


Hypoxia and other stress responses (e.g. ER stress response)

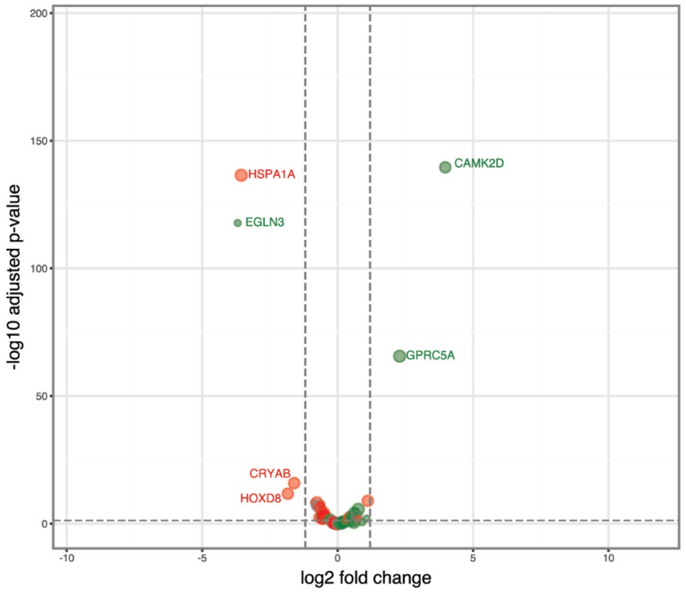


Inhibition of apoptotic signaling, downregulation of reactive oxygen species (ROS), and increased autophagy

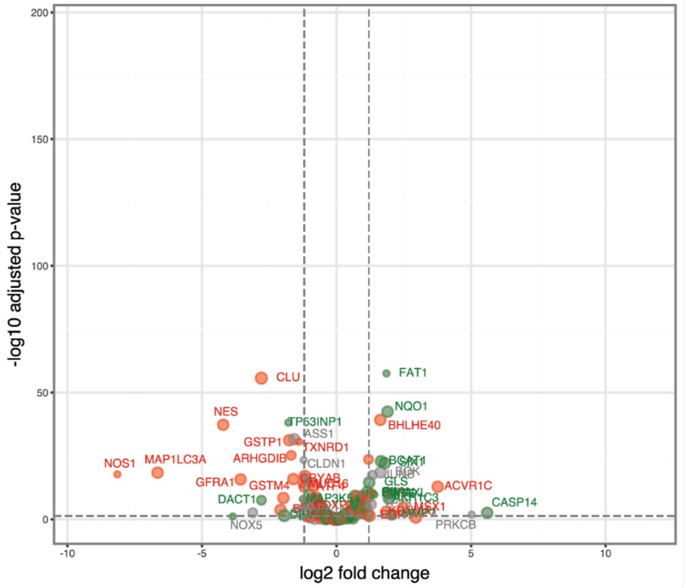

### Supplemental Figure 3 – OVCAR3 ResA (2 of 2)

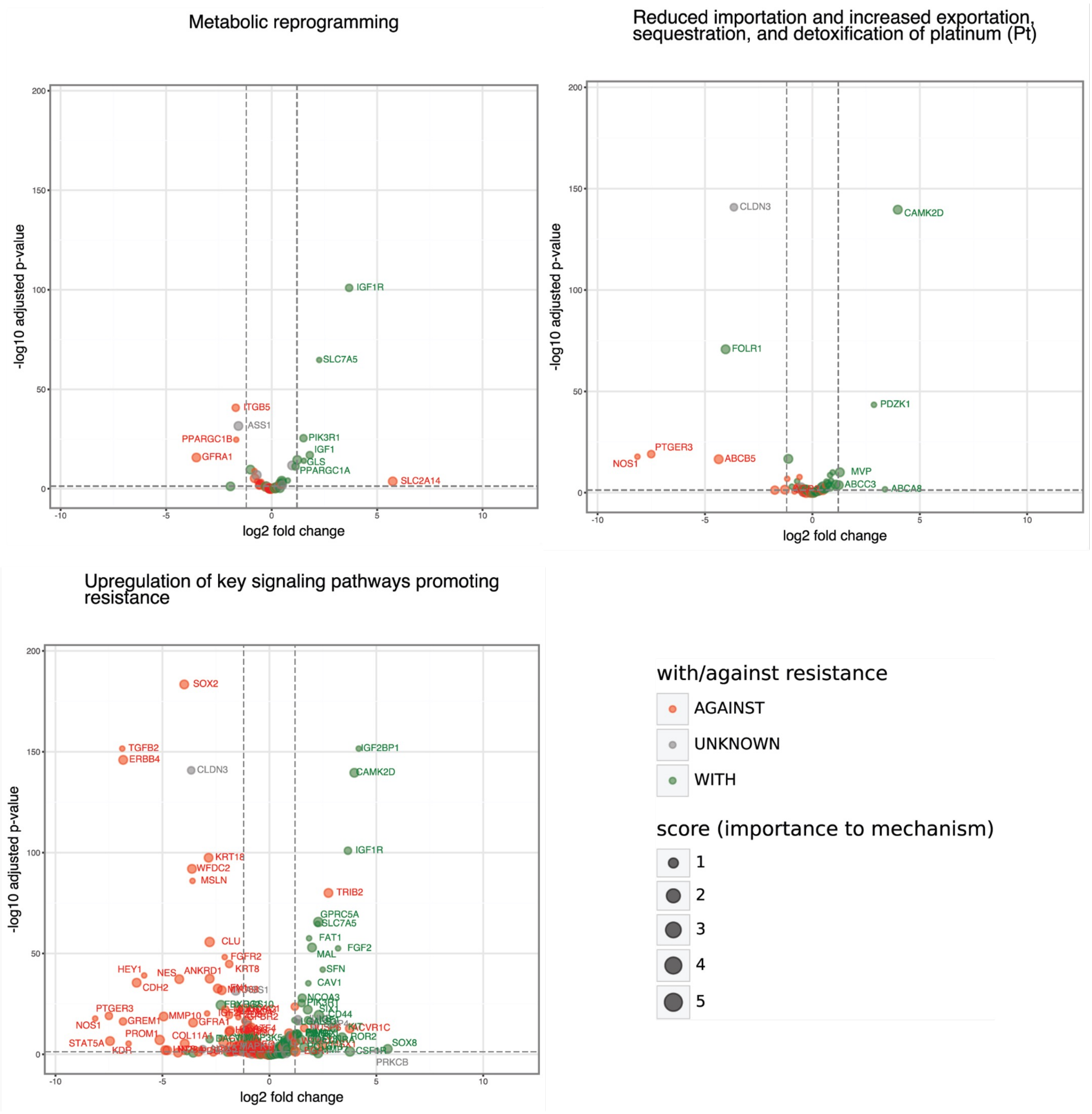

### Supplemental Figure 3 – OVCAR3 ResB (1 of 2)

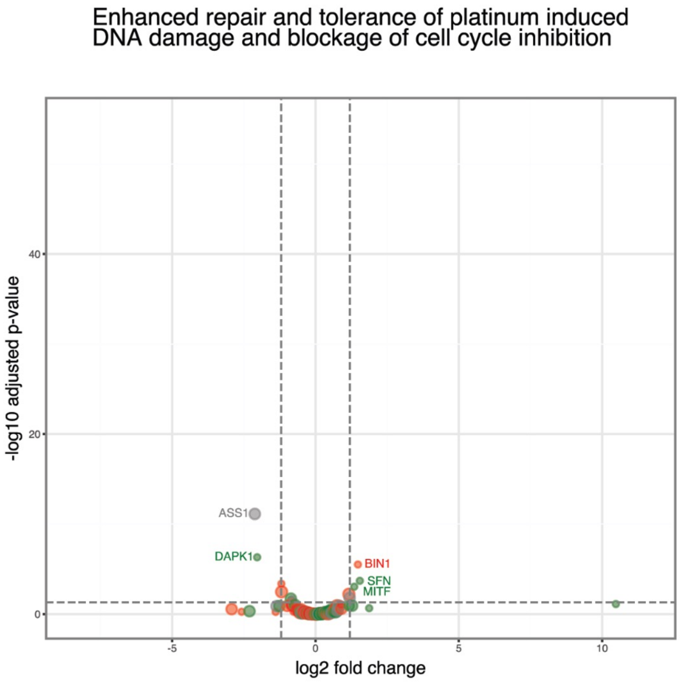


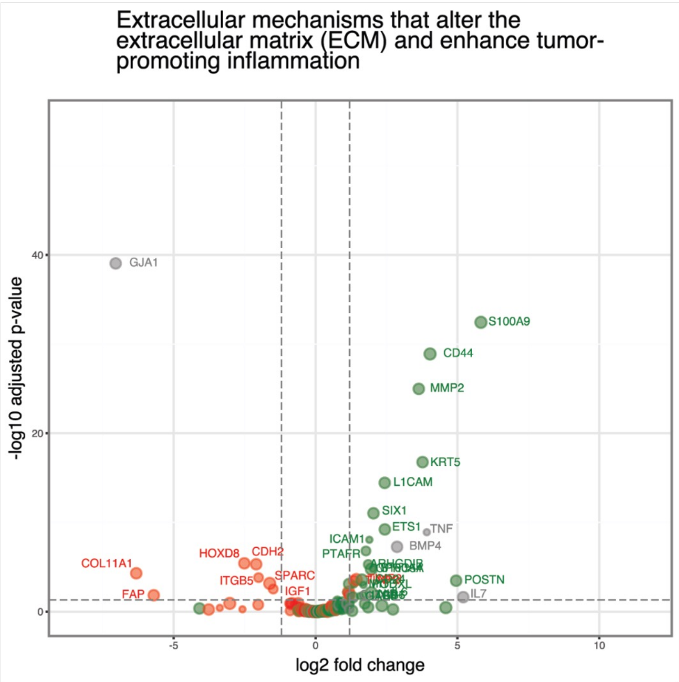


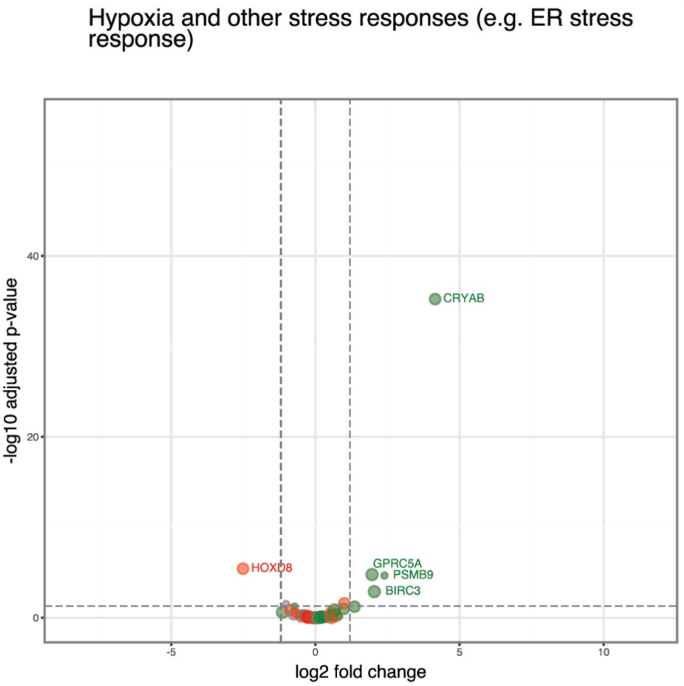


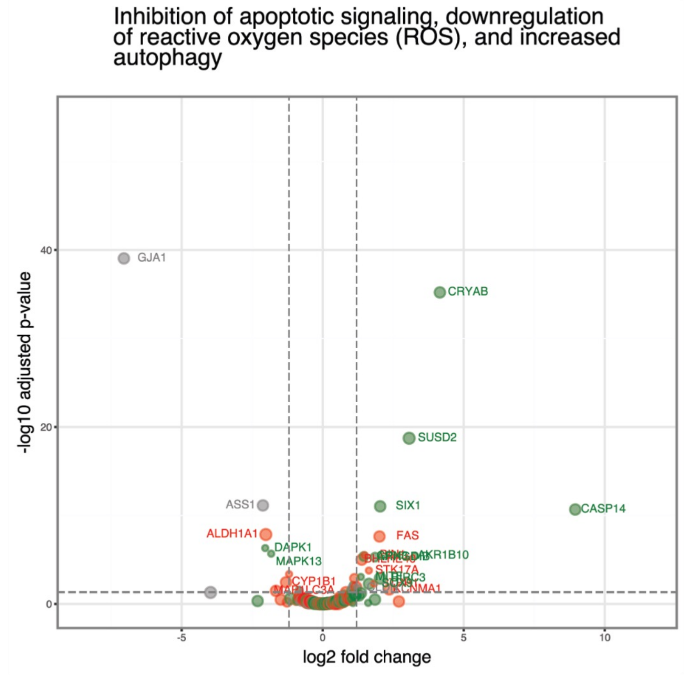

### Supplemental Figure 3 – OVCAR3 ResB (2 of 2)

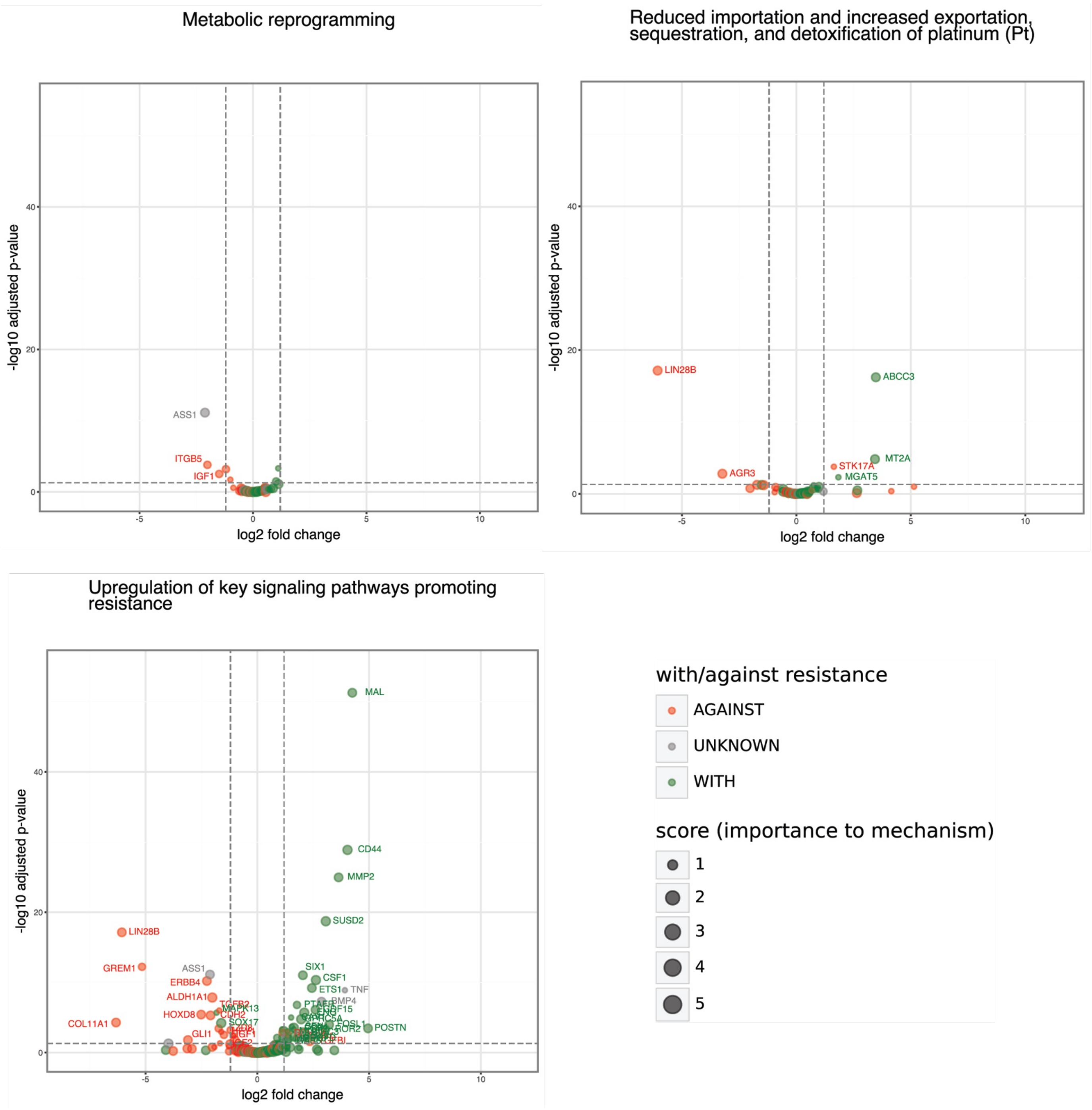

### Supplemental Figure 3 – OVCAR4 ResA (1 of 2)

Enhanced repair and tolerance of platinum induced DNA damage and blockage of cell cycle inhibition

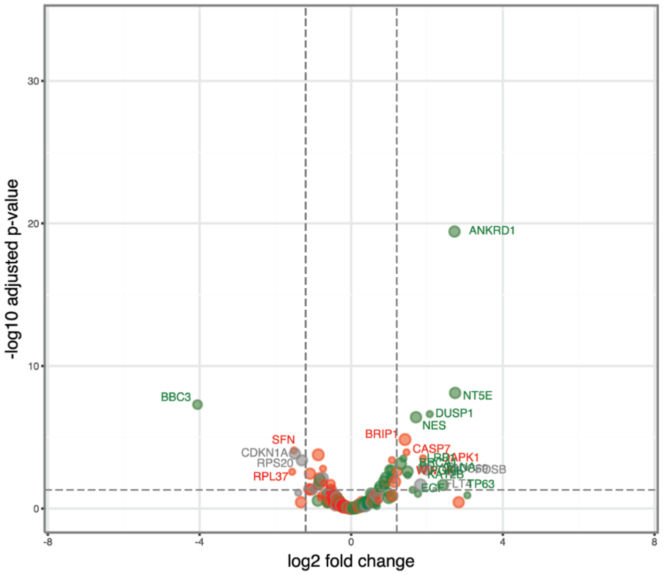


Extracellular mechanisms that alter the extracellular matrix (ECM) and enhance tumor-promoting inflammation

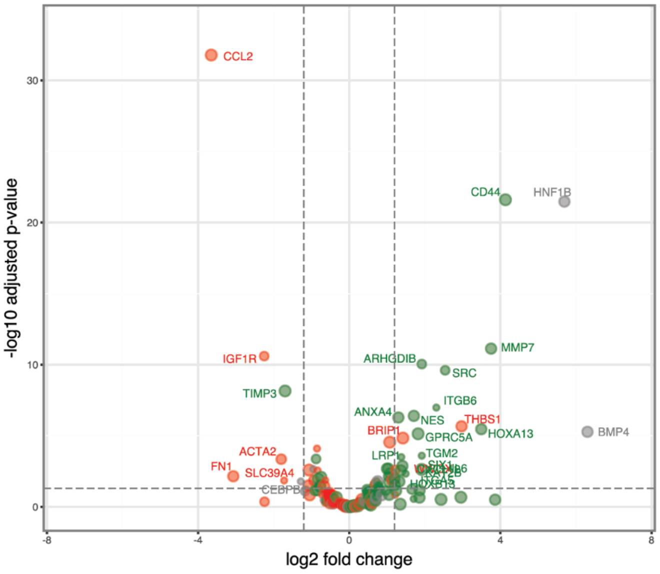


Hypoxia and other stress responses (e.g. ER stress response)

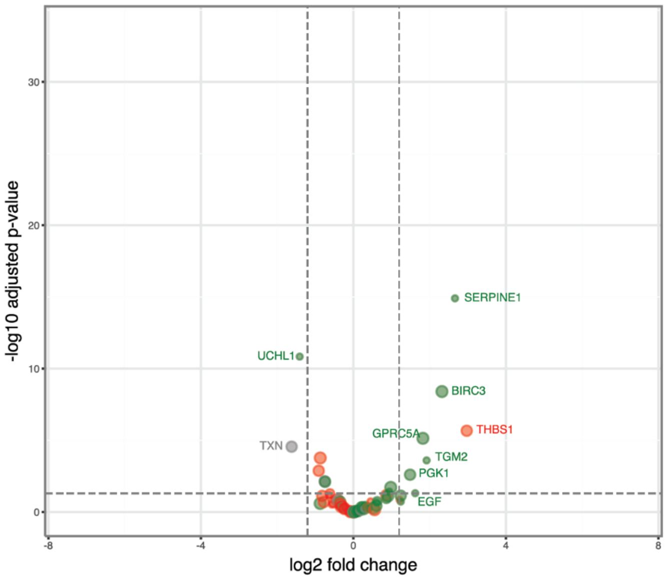


Inhibition of apoptotic signaling, downregulation of reactive oxygen species (ROS), and increased autophagy

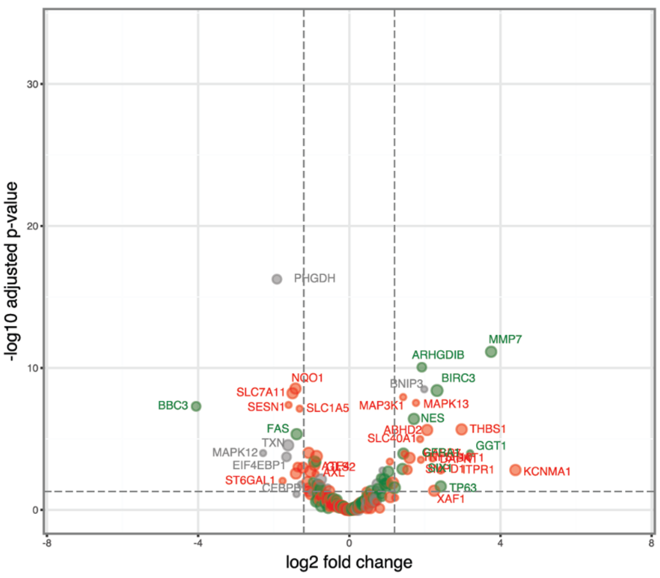

### Supplemental Figure 3 – OVCAR4 ResA (2 of 2)

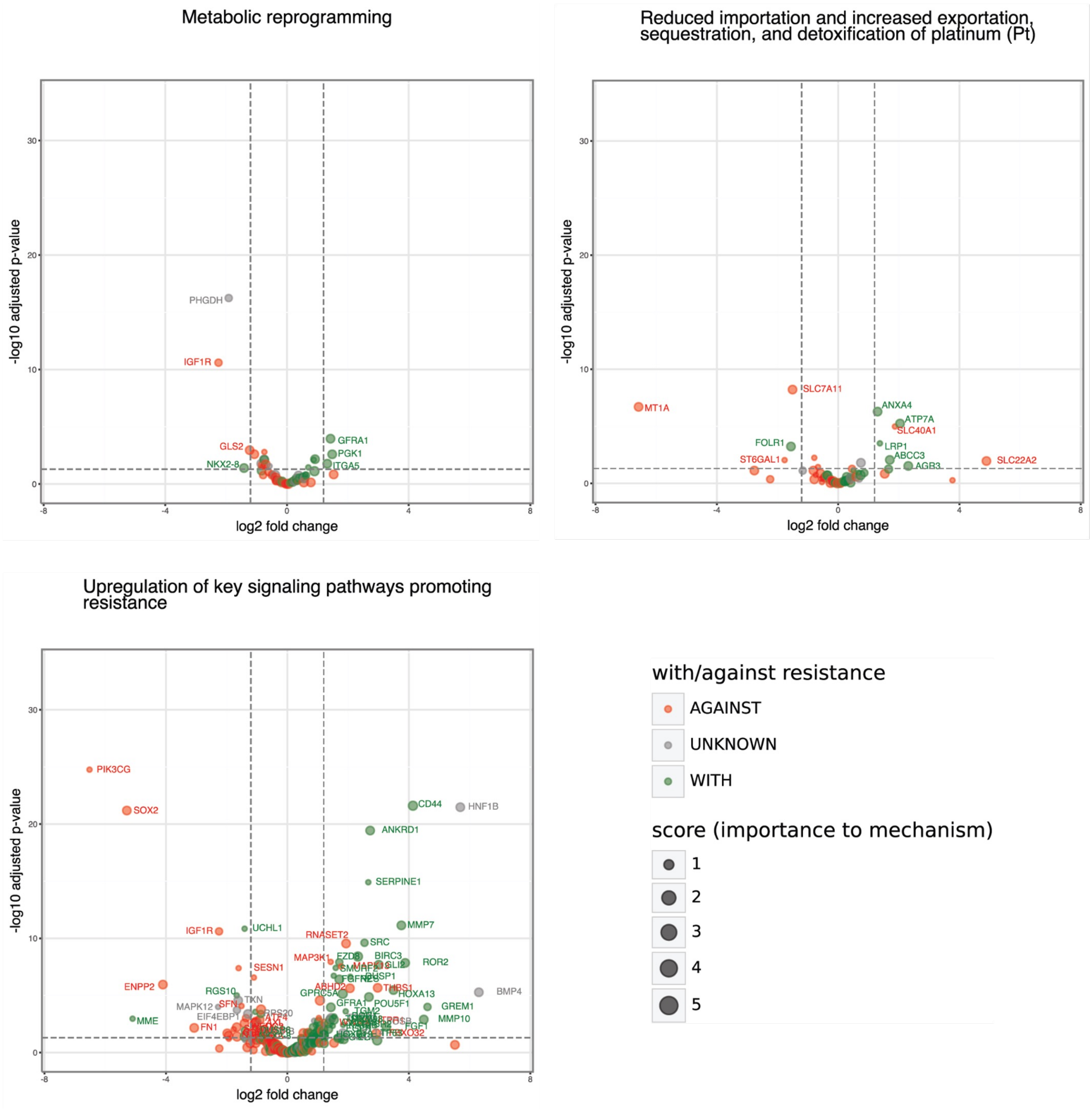

### Supplemental Figure 3 – OVCAR4 ResB (1 of 2)

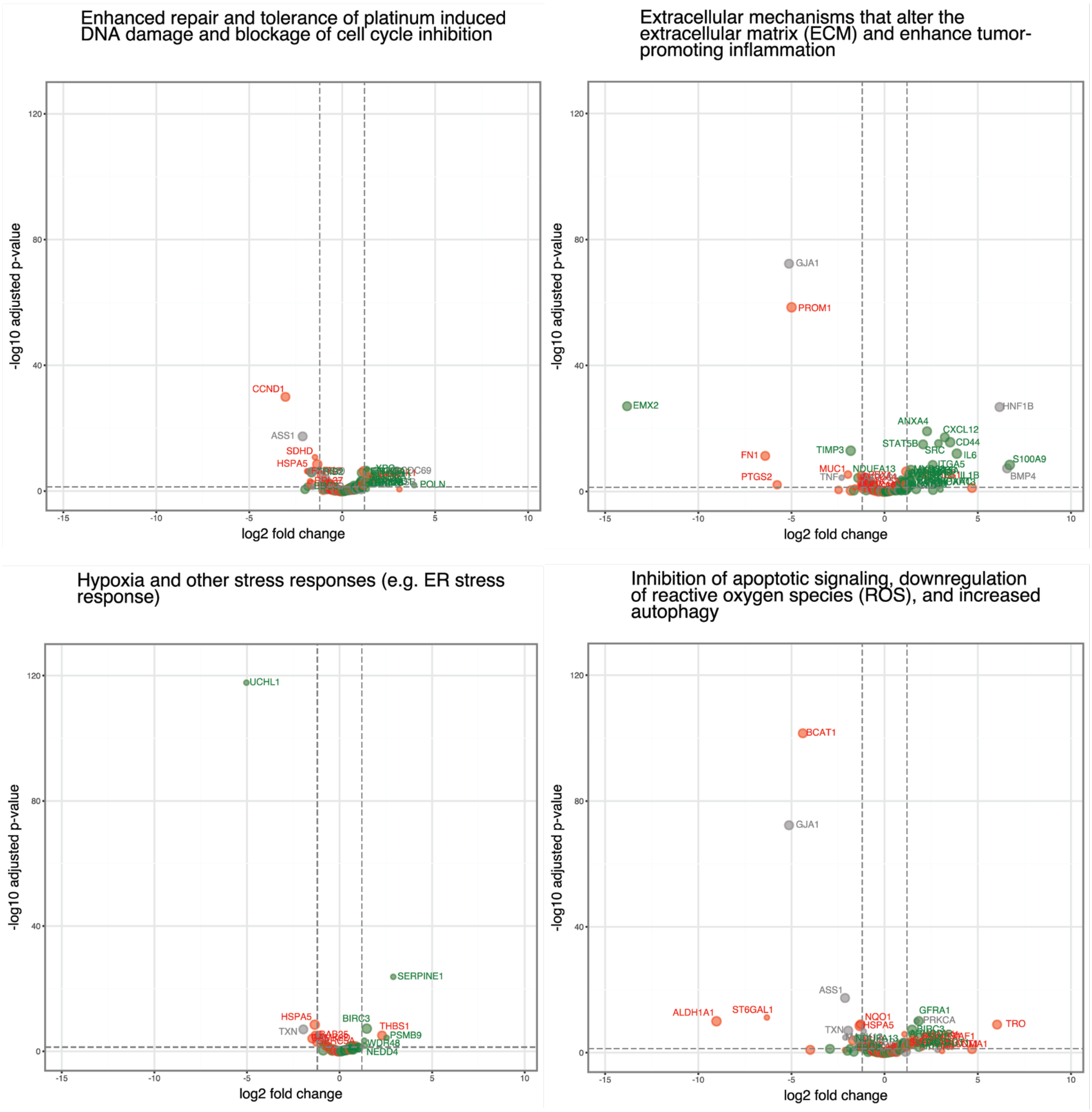

### Supplemental Figure 3 – OVCAR4 ResB (2 of 2)

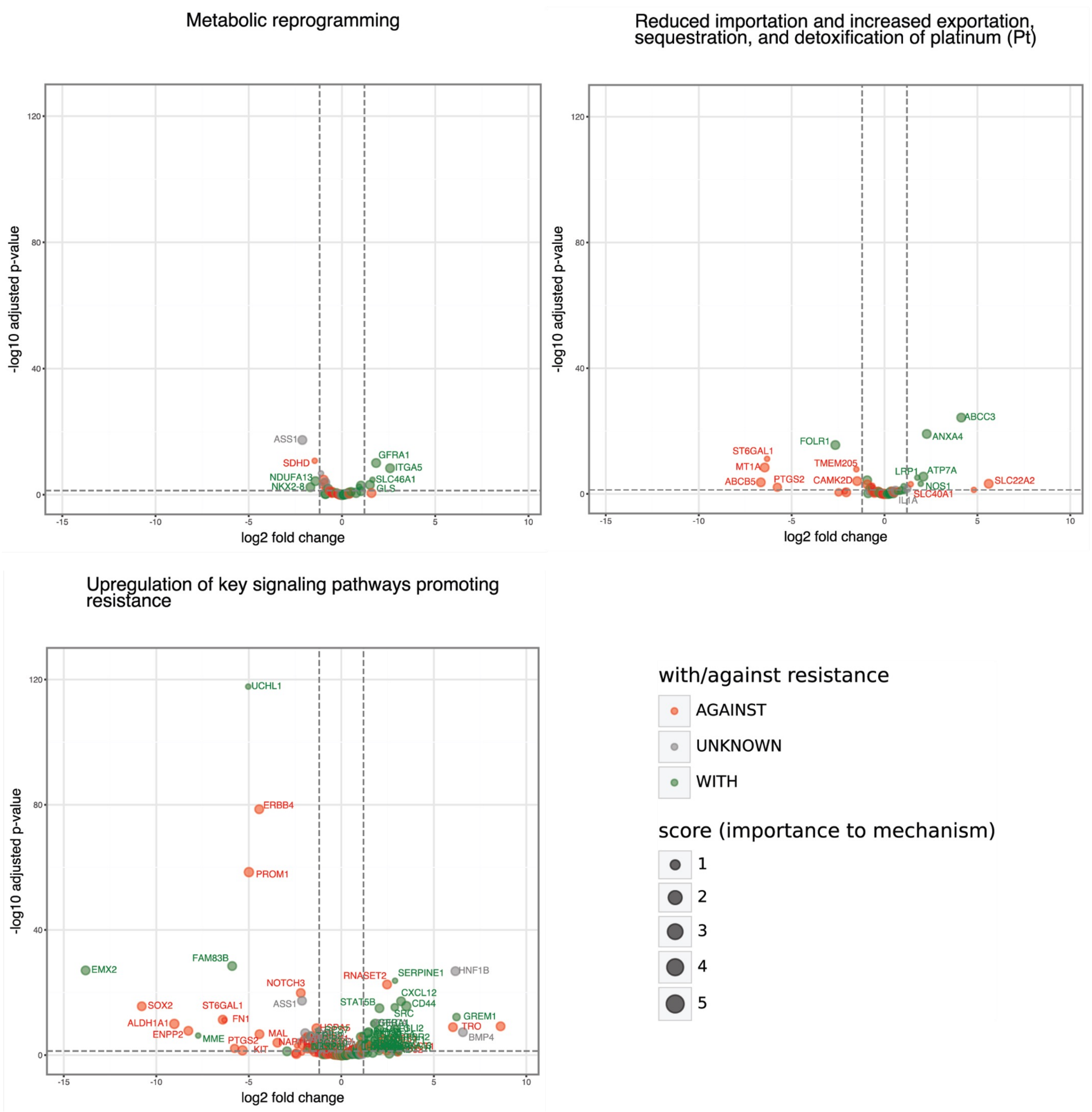

### Supplemental Figure 3 – PEO4 (1 of 2)

Enhanced repair and tolerance of platinum induced DNA damage and blockage of cell cycle inhibition

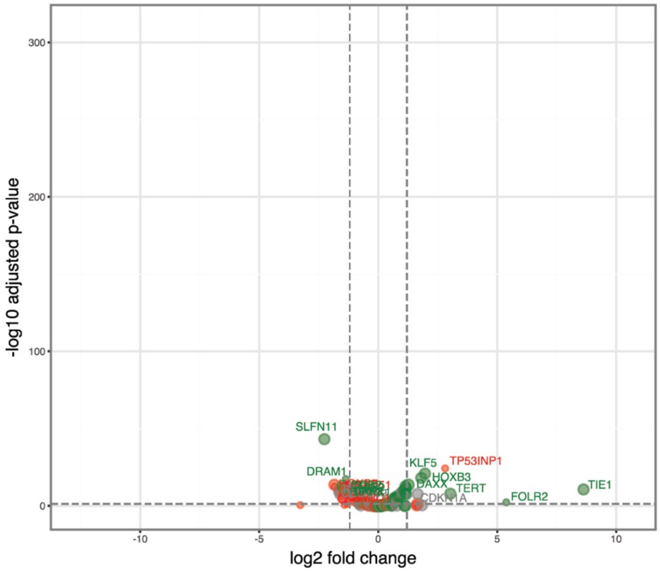


Extracellular mechanisms that alter the extracellular matrix (ECM) and enhance tumor-promoting inflammation

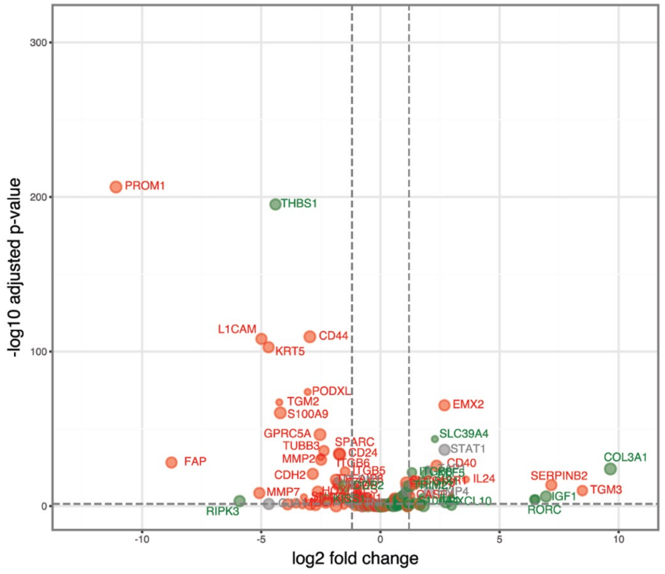


Hypoxia and other stress responses (e.g. ER stress response)

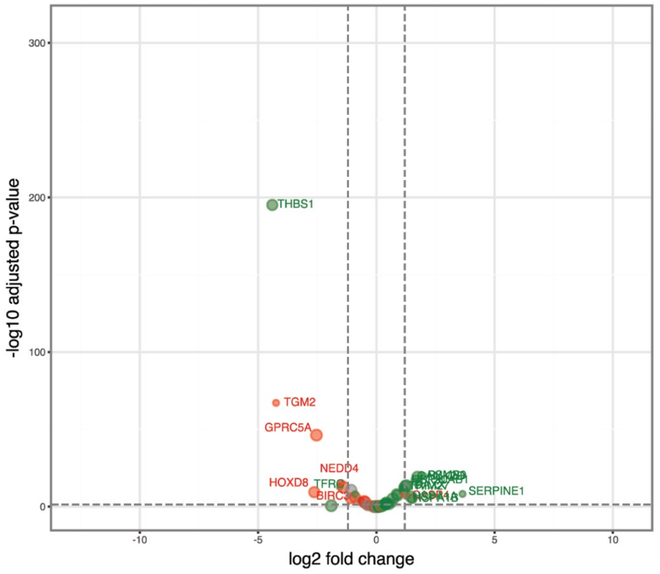


Inhibition of apoptotic signaling, downregulation of reactive oxygen species (ROS), and increased autophagy

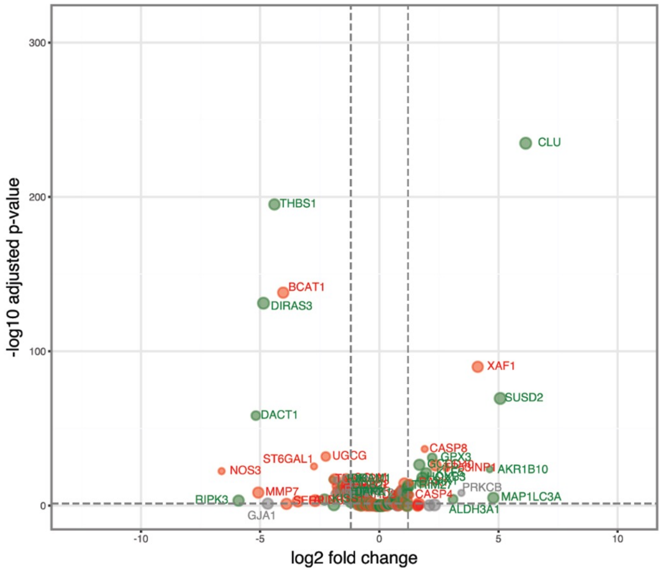

### Supplemental Figure 3– PEO4 (2 of 2)

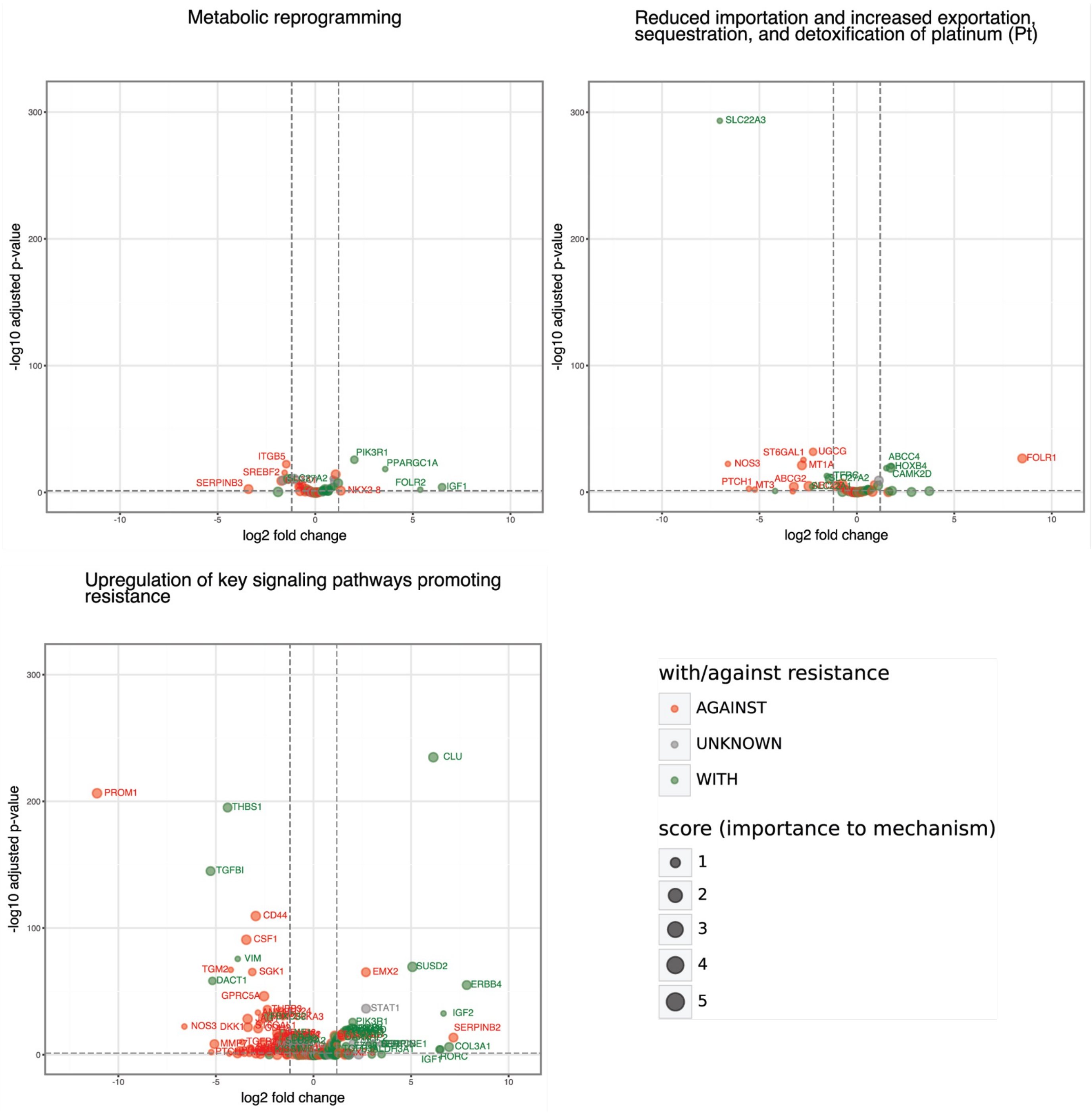

### Supplemental Figure 3 – PEO6 (1 of 2)

Enhanced repair and tolerance of platinum induced DNA damage and blockage of cell cycle inhibition

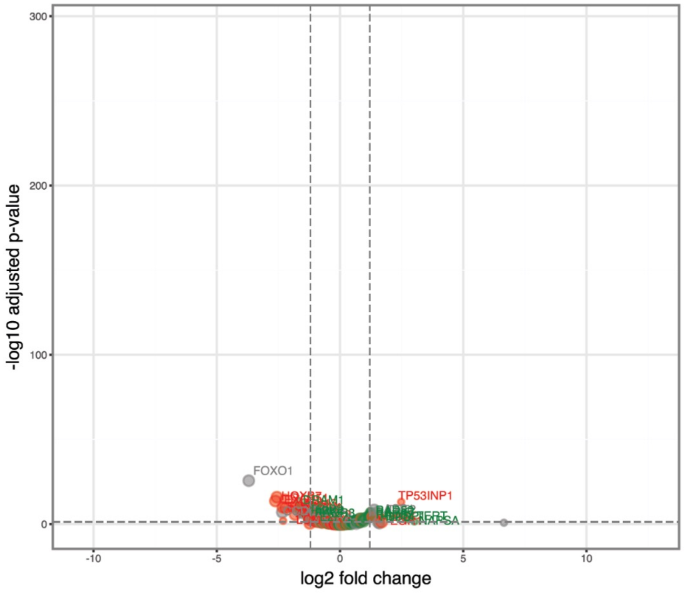


Extracellular mechanisms that alter the extracellular matrix (ECM) and enhance tumor-promoting inflammation

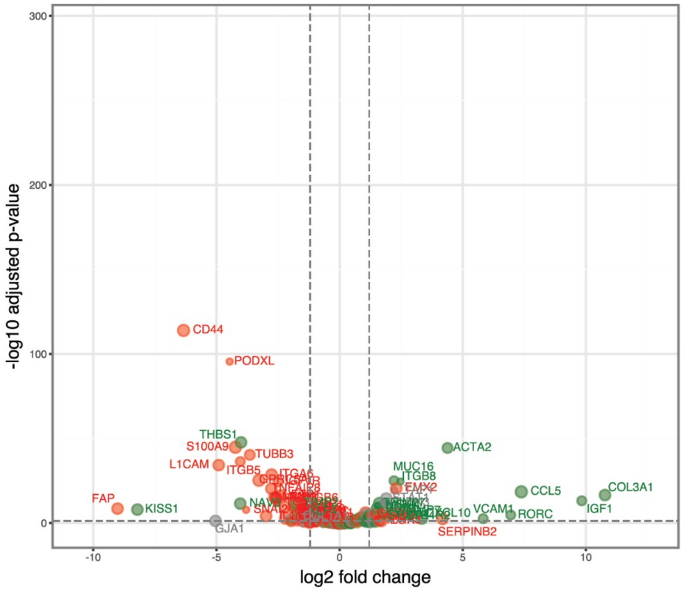


Hypoxia and other stress responses (e.g. ER stress response)

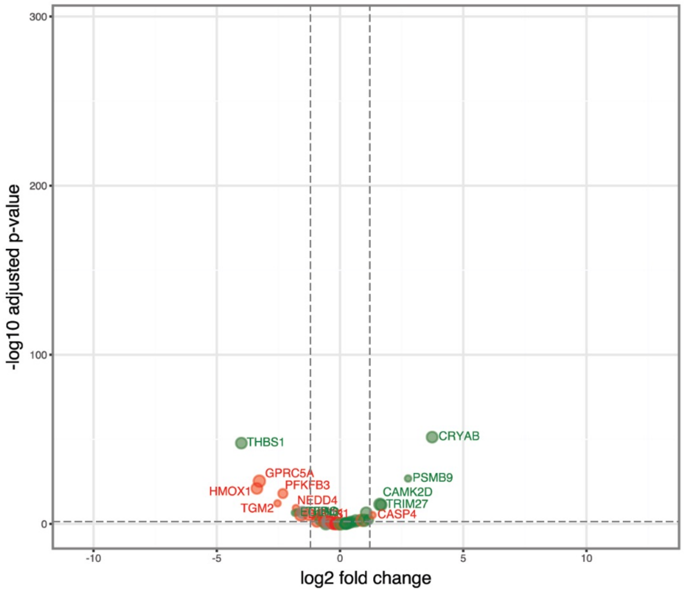


Inhibition of apoptotic signaling, downregulation of reactive oxygen species (ROS), and increased autophagy

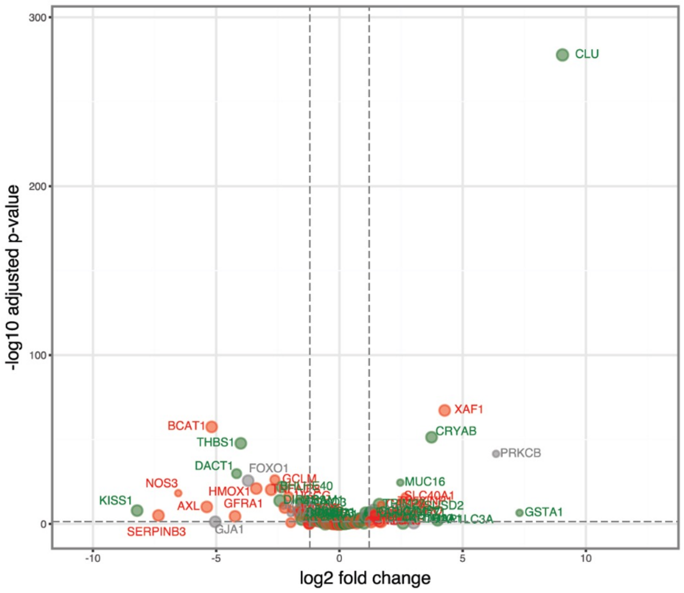

### Supplemental Figure 3 – PEO6 (2 of 2)

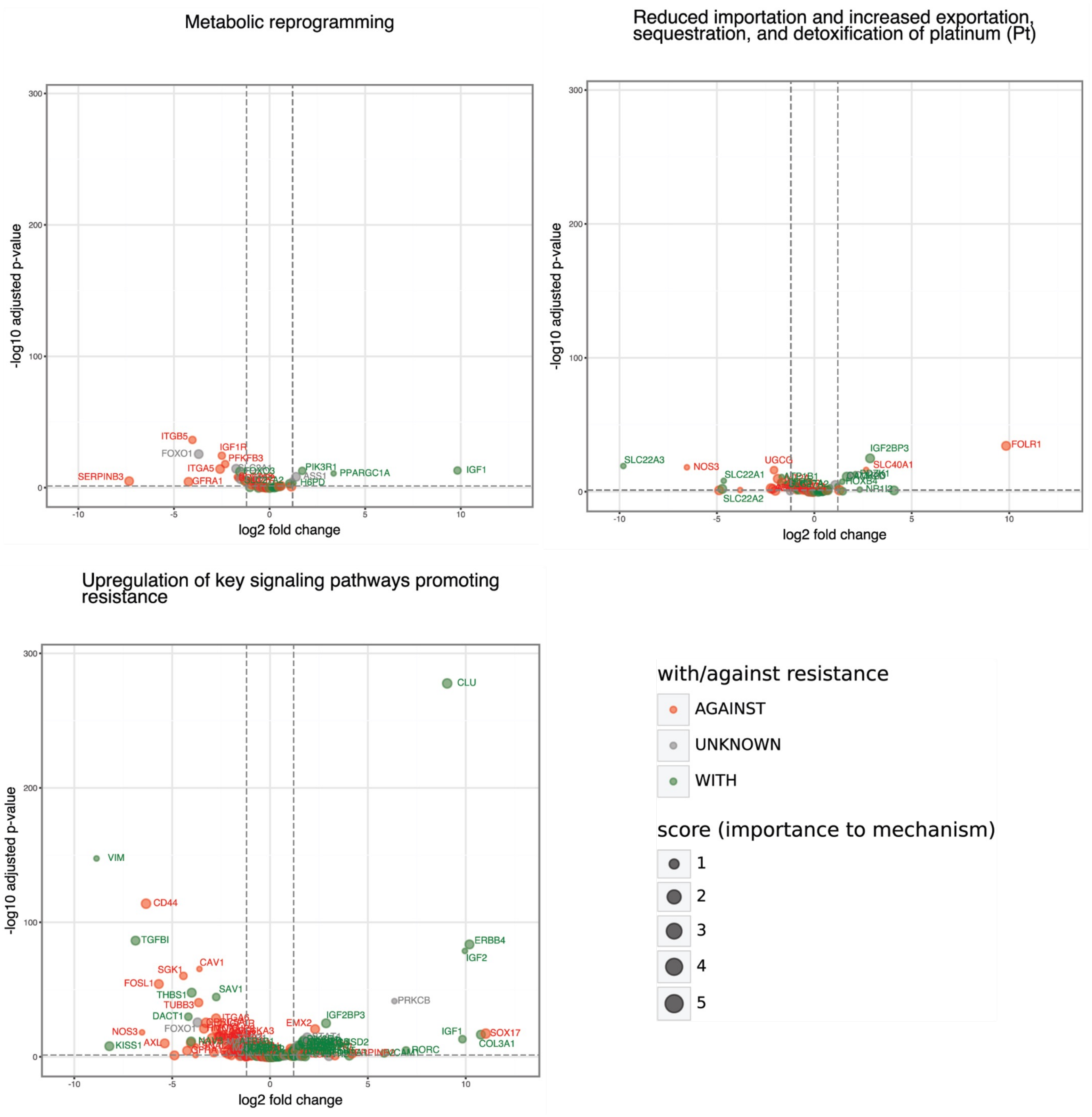

### Supplemental Figure 3 – PEA2 (1 of 2)

Enhanced repair and tolerance of platinum induced DNA damage and blockage of cell cycle inhibition

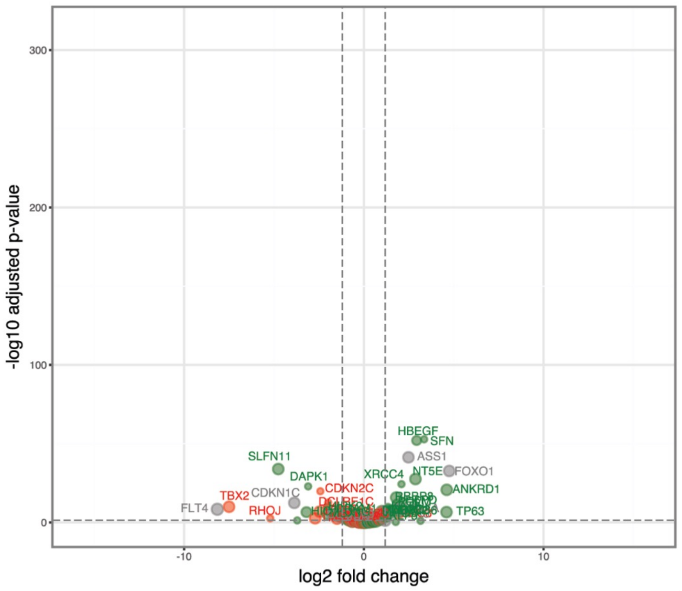


Extracellular mechanisms that alter the extracellular matrix (ECM) and enhance tumor-promoting inflammation

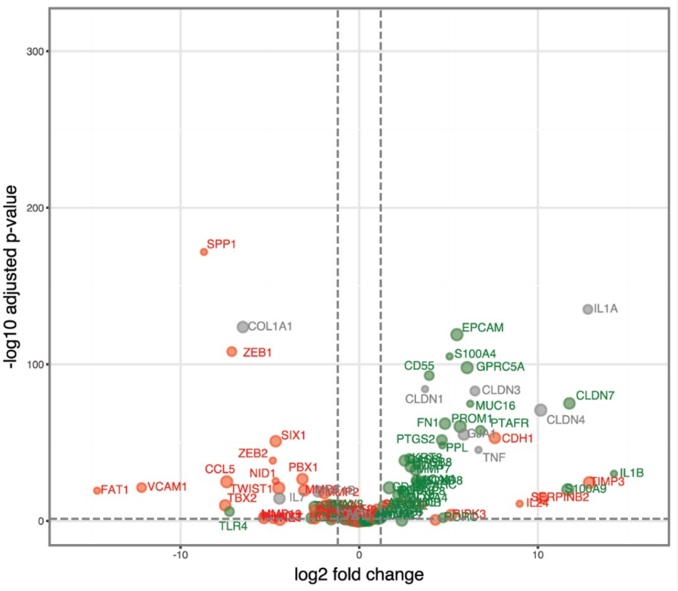


Hypoxia and other stress responses (e.g. ER stress response)

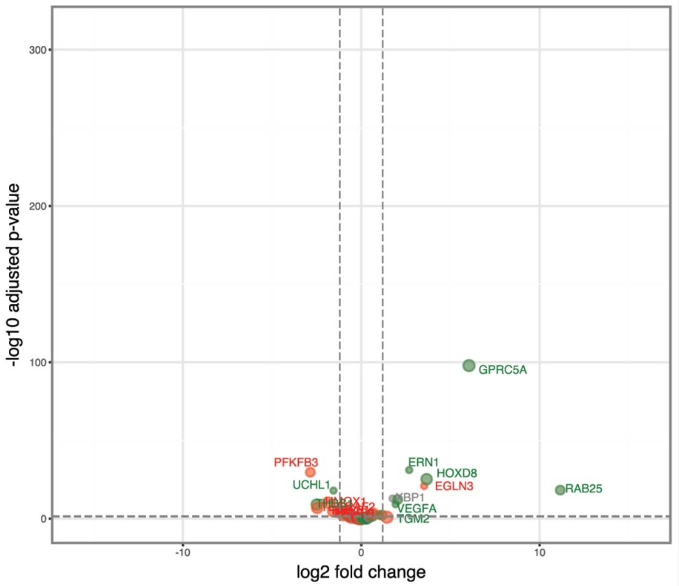


Inhibition of apoptotic signaling, downregulation of reactive oxygen species (ROS), and increased autophagy

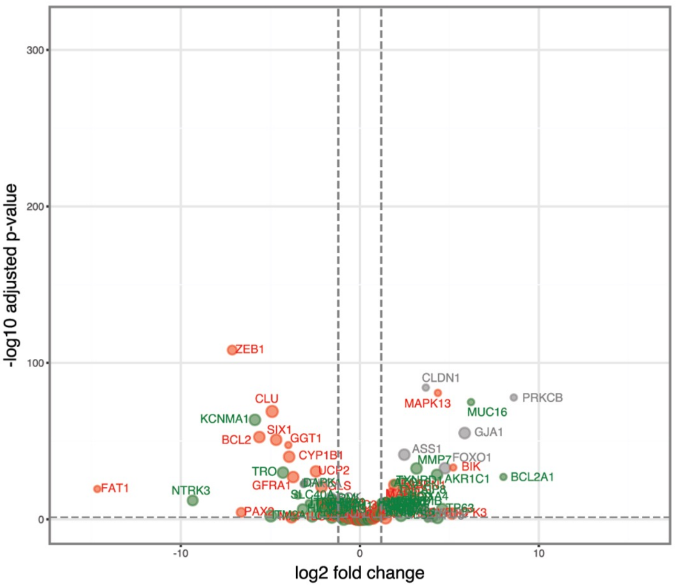

### Supplemental Figure 3 – PEA2 (2 of 2)

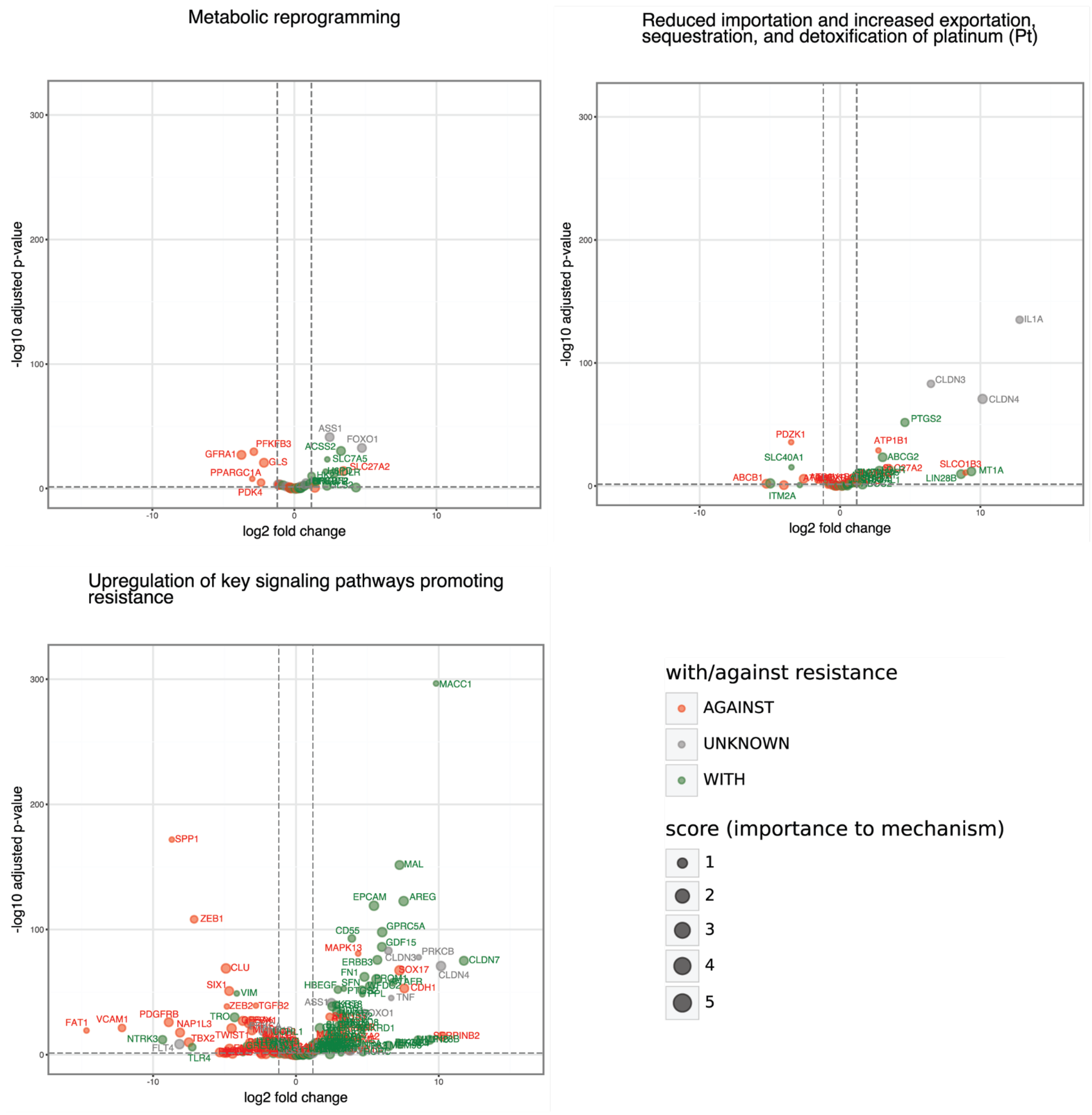

### Supplemental Figure 4

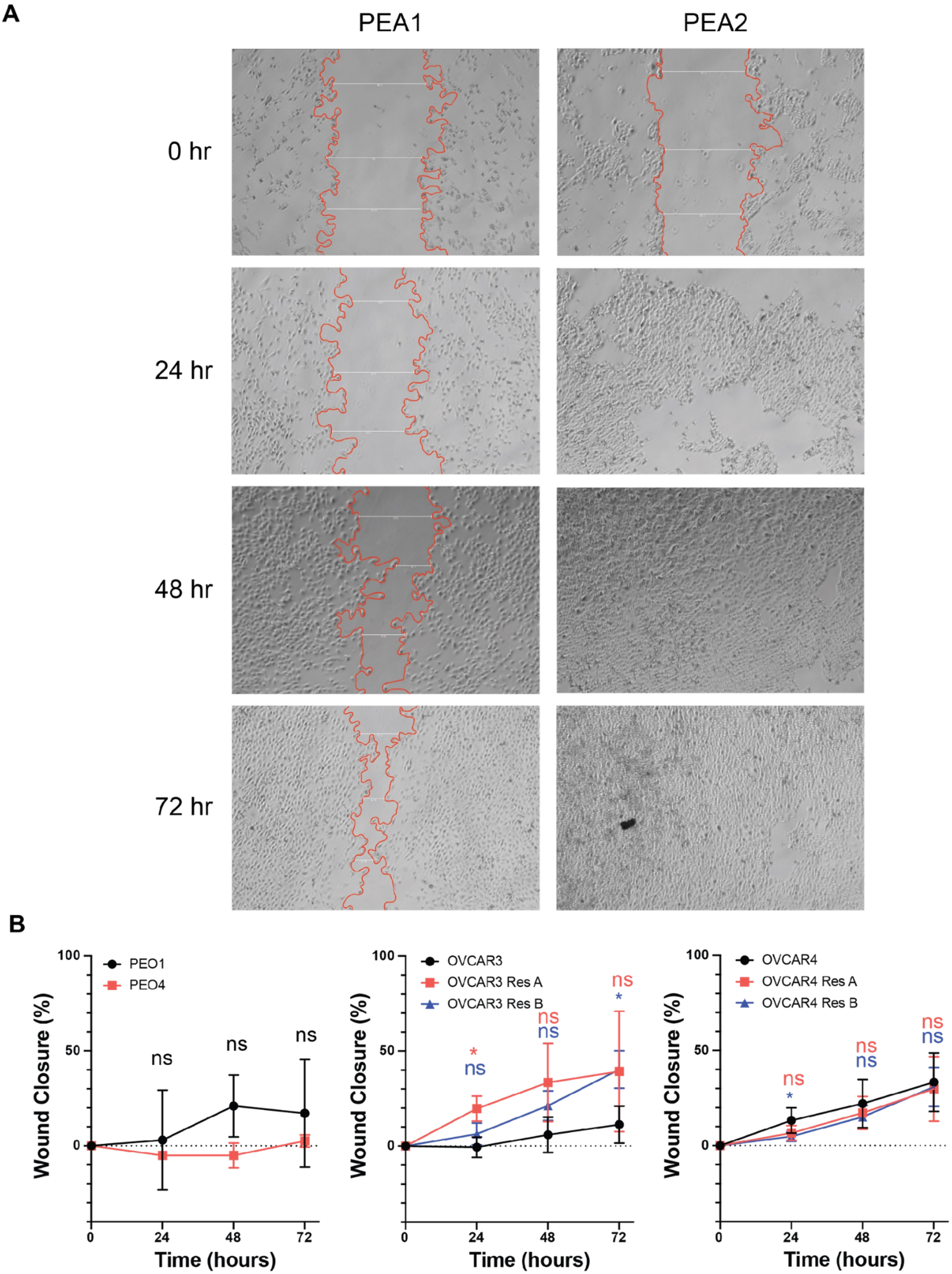

### Supplemental Figure 5

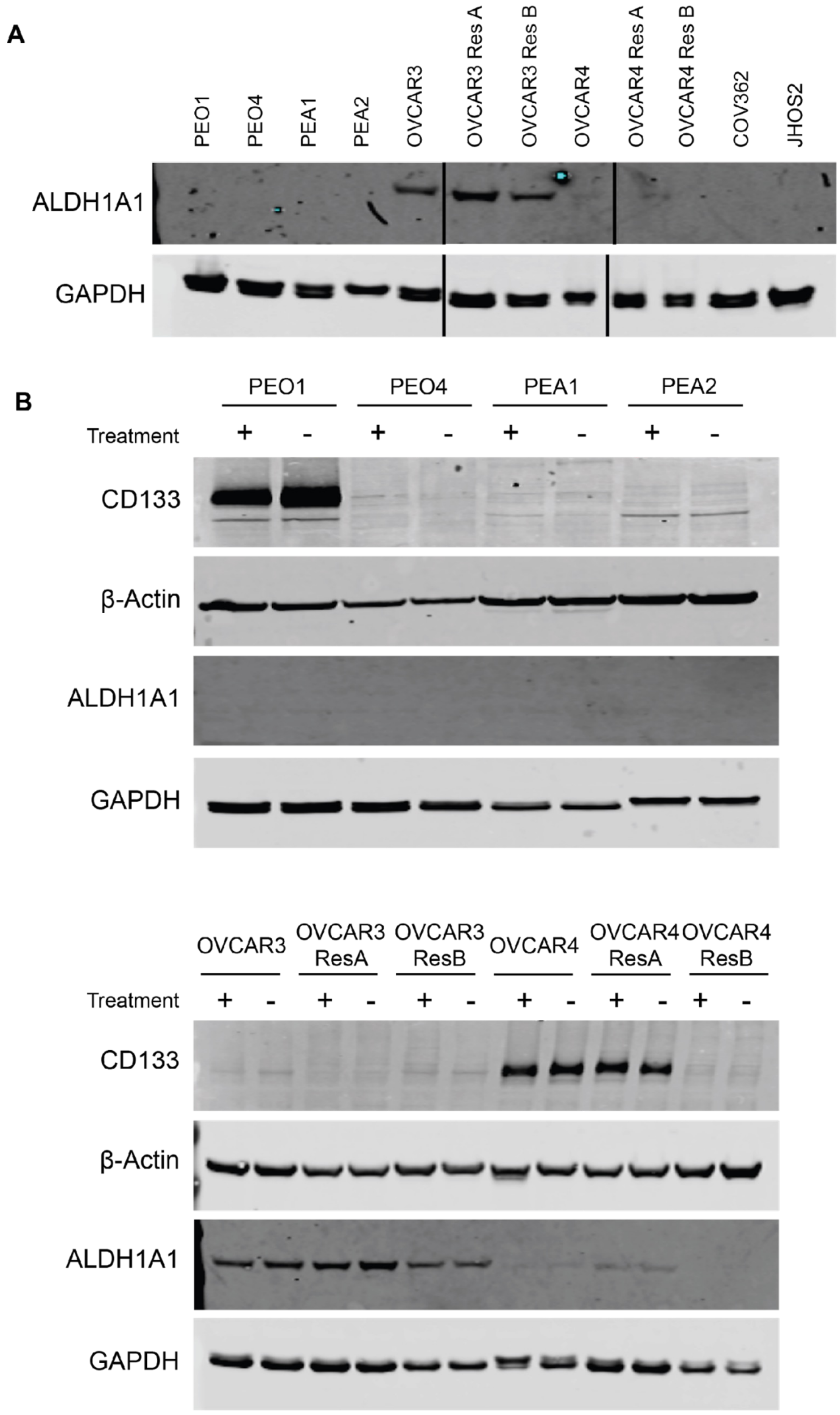

### Supplemental Figure 6A

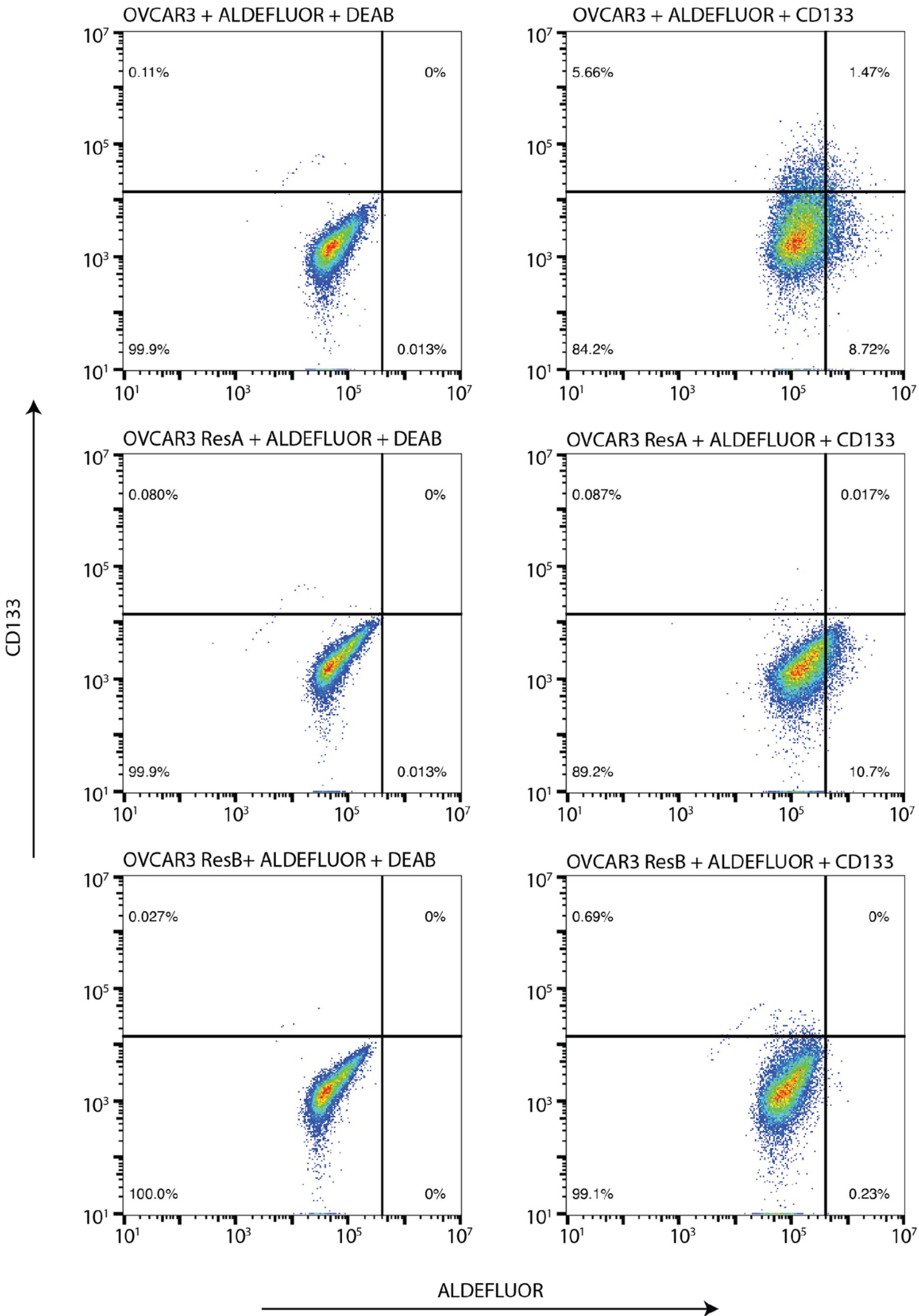

### Supplemental Figure 6B

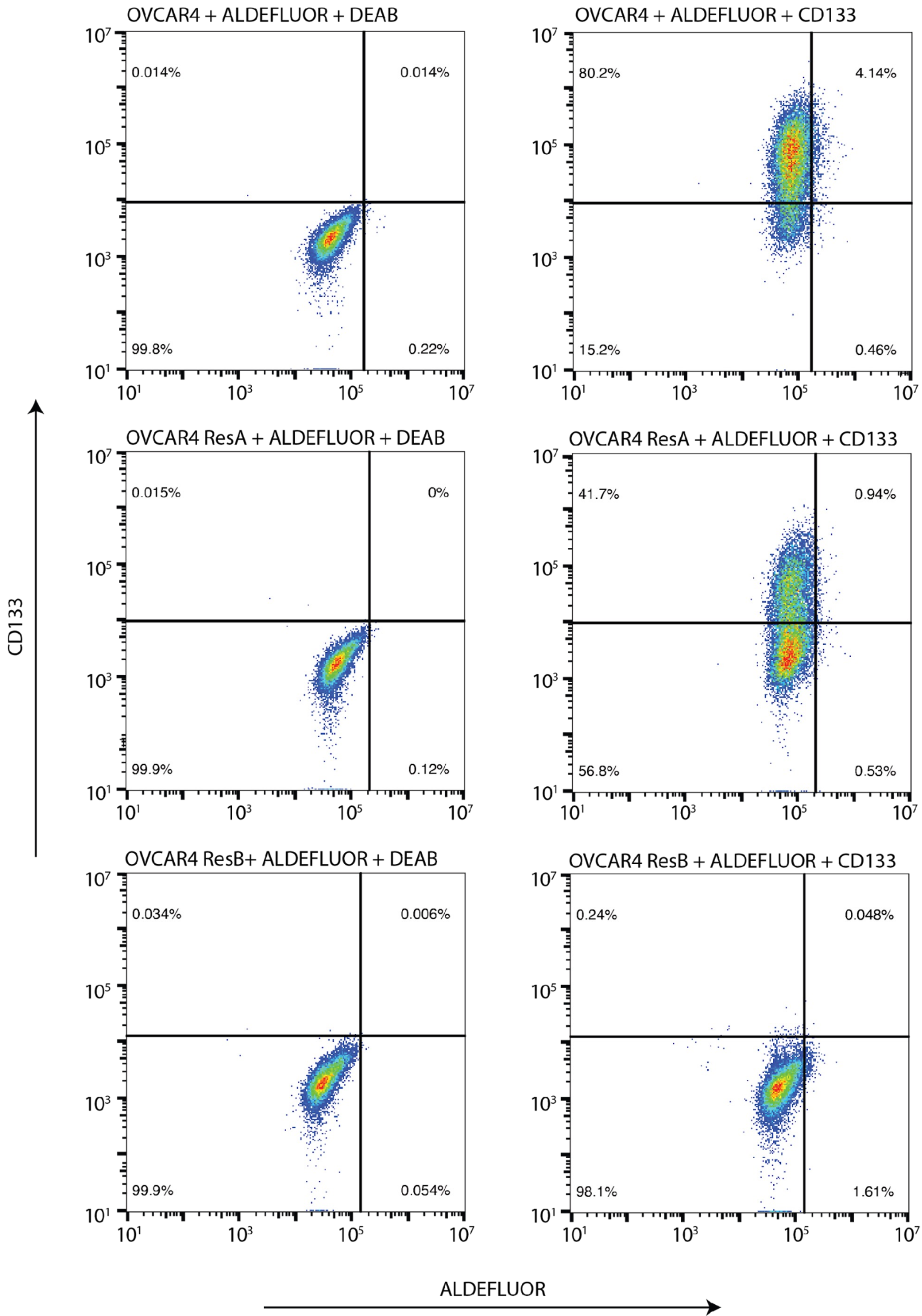

### Supplemental Figure 6C

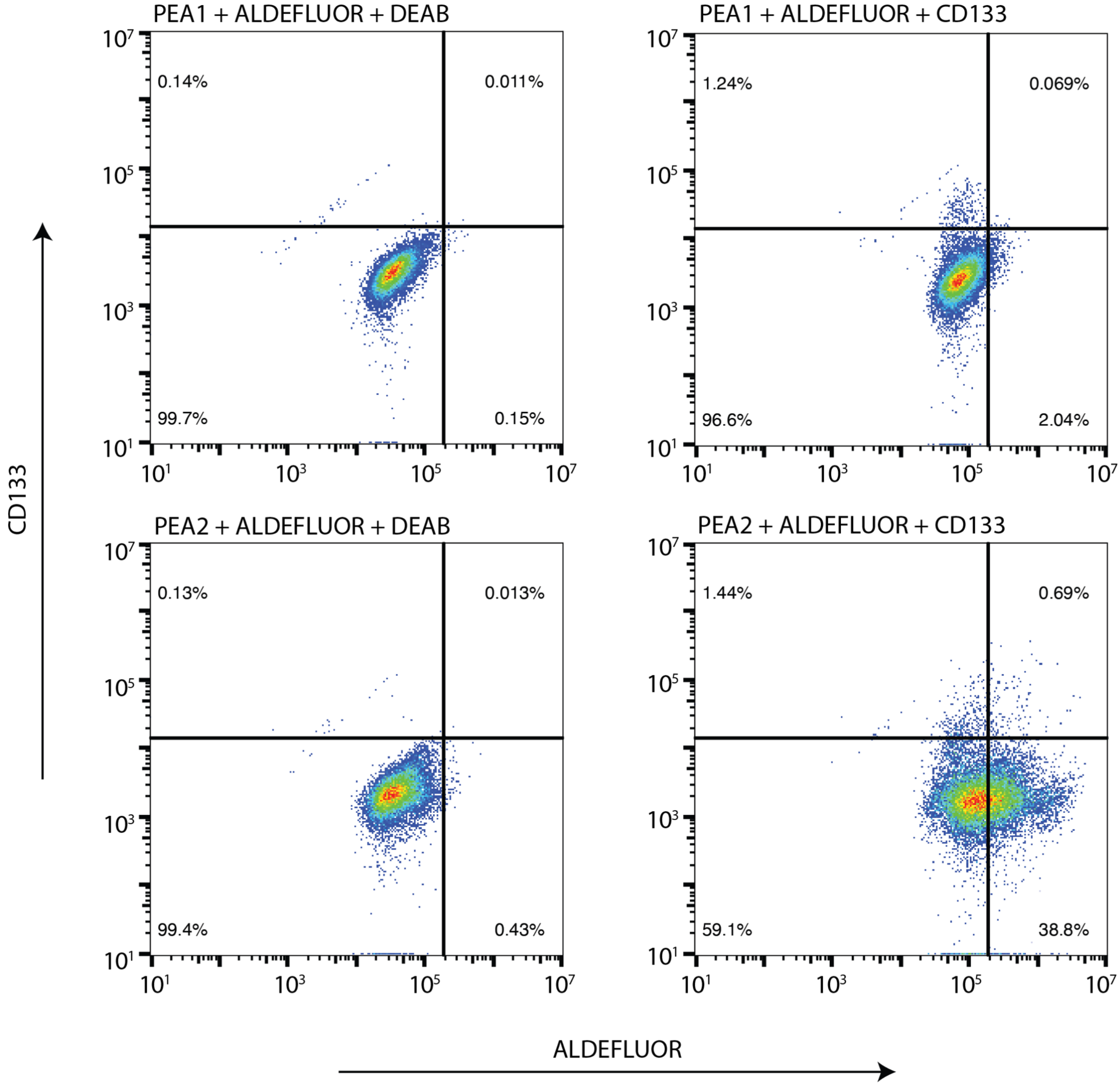

### Supplemental Figure 6D

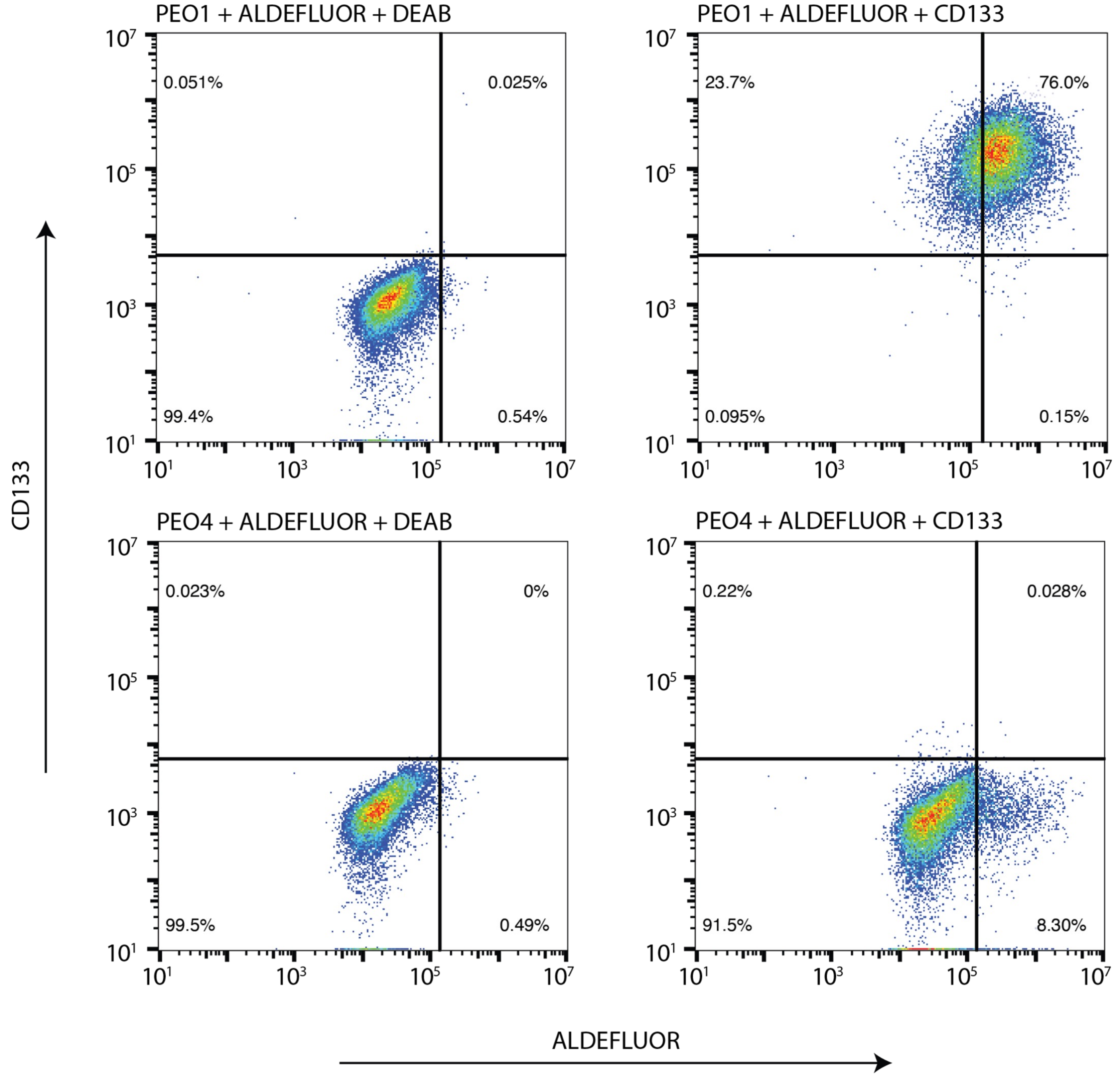
